# Psychotic-like experiences in children born very preterm: evidence from clinical and population-based cohorts

**DOI:** 10.64898/2026.08.13.744386

**Authors:** Claudia Aymerich, Marguerite Leoni, Alexandria O’Reilly Mescall, Zeyuan Sun, Divyangana Rakesh, Paola Dazzan, Emily Simonoff, A David Edwards, Lucy D. Vanes, Chiara Nosarti

## Abstract

**Background and aim:** Very preterm birth (VPT; ≤32 weeks’ gestation) is associated with an increased risk of later psychiatric disorders, including psychosis. Although psychosis typically emerges in adulthood, subclinical early signs along the psychosis continuum, such as psychotic-like experiences (PLEs), can be observed much earlier. We therefore aimed to study PLEs in childhood in VPT individuals recruited from a clinical cohort compared with full-term (FT) controls. We subsequently investigated whether findings could be replicated in an independent population-based cohort.

**Methods:** Primary analyses were conducted in the Brain, Immunity and Psychopathology (BIPP) study, including 197 children born VPT recruited through Neonatal Intensive Care Units and 72 FT controls assessed at a mean age of 10.50±1.77 years. Between-group differences in PLEs were then examined in the Adolescent Brain Cognitive Development (ABCD) study, including 149 children born VPT and 9519 FT controls assessed at a mean age of 9.94 ± 0.63 years. PLEs were assessed using the Prodromal Questionnaire-Brief Child Version (PQ-BC), yielding frequency and distress-related scores for both the total scale and three specific domains (unusual thought content, perceptual abnormalities, disorganised speech). Regression models tested associations between birth status (VPT and control) and PQ-BC scores adjusting for age, sex, and socio-economic status, with secondary models additionally adjusting for cognitive ability and broader psychopathology. Pooled analyses examined cohort effects (BIPP and ABCD) and cohort-by-group status (VPT and control) interactions.

**Results:** In BIPP, VPT birth was associated with higher PQ-BC total (β=1.61, p=0.004) and distress scores (β=0.78, p=0.039), with the strongest and most consistent associations observed for perceptual abnormalities across total score (sum of endorsed items), distressing items, and distress severity scores (all p≤0.01). These associations were attenuated but largely persisted after adjustment for cognitive ability and broader psychopathology, particularly for perceptual abnormalities. In ABCD, VPT birth was not significantly associated with global or domain-specific PQ-BC outcomes.

**Discussion:** VPT birth is associated with increased vulnerability to PLEs in childhood, particularly in the domain of perceptual abnormalities. The lack of clear replication in the population-based ABCD cohort may reflect differences in the composition of its VPT subgroup, which may not fully represent VPT individuals typically seen in clinical cohorts.

## 1. Introduction

Very preterm birth (VPT; ≤32 weeks of gestation) has been associated with an increased risk of mental health difficulties across development. Compared with individuals born at term, those born VPT are more likely to present neurodevelopmental and psychiatric problems, including attention-deficit / hyperactivity disorder (ADHD)^1^, emotional and behavioural difficulties^2^, and anxiety and mood-related symptoms^3^, among others.

Within this broader pattern of psychiatric vulnerability, increasing attention has been paid to psychosis-related outcomes. Epidemiological studies have suggested that individuals born preterm are at increased risk of later psychotic disorders^4^, and related perinatal factors such as low birth weight have also been associated with psychosis risk^5^. Together, these findings indicate that VPT birth may be linked not only to common psychiatric difficulties, but also to later liability for psychosis spectrum outcomes^6^.

Psychotic disorders are preceded, in a substantial proportion of cases, by prodromal or attenuated psychotic symptoms that may emerge years before the onset of a first psychotic episode^7,8^. These early subthreshold phenomena, commonly referred to as psychotic-like experiences (PLEs), include unusual perceptual experiences, beliefs, or thought content that resemble psychotic symptoms but do not meet the threshold for a psychotic disorder^9^. PLEs are relatively common in childhood and are often transient; however, when persistent, distressing, or associated with functional impairment, they may indicate increased vulnerability to subsequent psychotic disorder and broader psychopathology^10,11^. Their assessment in childhood is therefore potentially valuable for characterising early developmental pathways to mental-health difficulties. At the same time, PLEs can be difficult to interpret in children because unusual experiences and beliefs may partly reflect normative development, and their clinical significance depends on factors such as frequency, persistence, and associated distress^12^.

At present, it remains unclear whether individuals born VPT already show an increased risk of PLEs during childhood. Addressing this question is important because childhood PLEs may represent an early transdiagnostic marker of vulnerability, being associated not only with later psychotic disorders but also with broader emotional, behavioural, and psychiatric difficulties^13^. It is also unknown whether this potential vulnerability is expressed broadly across symptom domains or is more specifically characterised by certain experiences. Importantly, investigating PLEs in children born VPT also requires consideration of cognitive functioning and broader psychopathology, to determine whether elevated PLEs reflect a specific psychosis-related vulnerability or are better understood as part of more general neurodevelopmental difficulties. This is particularly relevant given that cognitive deficits are widely considered a core feature of psychosis, evident not only from the first psychotic episode^14^ but also during earlier stages of illness^15^.

This study had two main objectives. First, we aimed to investigate whether VPT birth is associated with an increased risk of PLEs during childhood in a birth cohort of children who were born VPT in the UK, to characterise the profile of these symptoms, and to determine whether increased PLEs represent an independent effect of VPT birth or are better explained by comorbid cognitive and psychiatric difficulties. Second, we sought to replicate and validate these findings in an independent population-based cohort of children born in the US. Examining this question across cohorts with contrasting recruitment designs provides an opportunity to assess the robustness and generalisability of any observed associations.

## 2. Methods

### 2.1 Study design and participants

#### This study used data from two cohorts

The Brain, Immunity and Psychopathology (BIPP) study recruited VPT children born between 2010 and 2013 in hospitals within the North and Southwest London Perinatal Networks and their parents who had previously participated in the Evaluation of Preterm Imaging study (ePrime; EudraCT 2009-011602-42^16^). Infants enrolled in ePrime were eligible if they were born before 33 weeks’ gestation, their mothers were older than 16 years, and their mothers were not hospital inpatients at the time of recruitment. Exclusion criteria included major congenital malformation, contraindications to magnetic resonance imaging, parents unable to speak English, or involvement in child protection proceedings^16^. VPT participants underwent neurodevelopmental assessments at around age two and four^17,18^. The current study focuses on the follow-up assessment conducted between the ages of 8 and 11 years (BIPP study), which further recruited a control group of age- and sex-matched participants born at term (i.e., after 37 completed weeks of gestation) from the local community^2^. In addition to the ePRIME exclusion criteria, BIPP excluded participants who were not attending a mainstream school. The final analytic sample comprised 197 children born VPT and 72 control children born full term (FT).

The Adolescent Brain Cognitive Development (ABCD) study is an ongoing nationwide prospective observational study of brain development that follows 11,875 participants from ages 9- 11 into early adulthood across 21 sites across the United States. It was designed to capture a diverse range of geographic, socio-economic, ethnic, and health backgrounds^19,20^. At all sites, parents provided written informed consent and children provided assent, with all procedures conducted in accordance with Institutional Review Board guidelines. For the present study, data were drawn from the baseline cohort included in ABCD Annual Curated Data Release 3.0 (https://abcdstudy.org/), comprising children aged 9-11 years of age. Because ABCD is a population-based cohort recruited in mid-childhood rather than at birth, it was used here as an independent validation sample rather than as a directly equivalent replication cohort. The final analytic sample comprised 149 children born VPT and 9,519 children born FT.

### 2.2 Definition of preterm birth

In BIPP, gestational age at birth for VPT participants was collected from neonatal medical discharge notes. For controls, gestational age was retrospectively collected from their parents. All control participants were born at or after 37 weeks’ gestation (i.e., FT).

In ABCD, VPT status was determined using two parent-reported questions: “Was the child born prematurely?” and “About how many weeks premature was the child when they were born?”. Children were classified as FT if parents responded “no” to the first question, and as VPT if parents responded “yes” and reported that the child had been born 8 or more weeks premature, corresponding to a gestational age of 32 weeks or less. However, unlike BIPP, ABCD excluded children born extremely preterm (<28 weeks’ gestation)^21^. To maximize comparability across cohorts, participants born between 33- and 37-weeks’ gestation were excluded from all analyses.

### 2.3 Socio-demographic measures

Sociodemographic variables available in both cohorts included sex, ethnicity (self-reported), age at assessment, and highest educational attainment of the primary caregiver. Socio-economic status was indexed using area-level measures of neighbourhood deprivation. In BIPP, neighbourhood deprivation was measured using the 2019 Index of Multiple Deprivation (IMD) decile derived from the maternal postcode at recruitment, with higher deciles indicating lower deprivation^22^. In ABCD, neighbourhood deprivation was measured using the scaled weighted Area Deprivation Index (ADI)^23^ score derived from the participant’s primary residential address at baseline, with higher scores indicating greater deprivation. Caregiver educational attainment was used to characterise the samples descriptively and was not included as a covariate in the regression analyses.

### 2.4 Mental health assessments

In BIPP, dimensional psychopathology was indexed using the internalising, externalising, and total difficulties scores of the Strengths and Difficulties Questionnaire (SDQ)^24^. Internalising difficulties were derived from the emotional symptoms and peer problems subscales, whereas externalising difficulties were derived from the conduct problems and hyperactivity/inattention subscales^25^. In ABCD, dimensional psychopathology was assessed using the parent-reported Child Behaviour Checklist (CBCL)^26^. The Internalising Problems, Externalising Problems, and Total Problems scores were used in the analyses. Prior work has shown that SDQ and CBCL assess broadly comparable dimensions of child psychopathology and support score linking across internalising, externalising, and total problem domains^27^.

PLEs were assessed in both cohorts using the validated Prodromal Questionnaire – Brief Child Version (PQ-BC)^28^. The PQ-BC is a 21-item self-report measure that assesses prodromal symptoms and includes a visual response Likert-type scale to rate associated distress. In the present study, we examined the Total score (sum of endorsed items), Distressing Items Score (sum of endorsed items also rated as distressing), and the Distress Severity Score (sum of distress ratings across all endorsed PLEs).

In addition, PQ-BC items were classified into three symptom domains based on the structure of the Structured Interview for Psychosis-Risk Symptoms (SIPS)^29^, against which the PQ-B was originally validated^30^. Accordingly, three symptom domains were defined: unusual thought content, perceptual abnormalities, and disorganised speech. The items included in each domain are listed in eMethods 1. Total scores, Distressing Items Scores, and Distress Severity Scores were derived for each of these domains separately.

### 2.5 Cognitive assessments

In BIPP, cognitive ability was measured using the age-adjusted full-scale composite score of the Wechsler Intelligence Scale for Children (WISC). In ABCD, cognitive ability was assessed using the age-normed scaled score from the Matrix Reasoning subtest of the WISC-V^31^. Higher scores indicate better cognitive performance.

### 2.6 Statistical analysis

All analyses were conducted in R version 4.2.2^32^. The primary analyses were conducted in the BIPP cohort. Secondary analyses were then performed in the ABCD cohort to assess whether the pattern of findings could be reproduced in an independent sample.

To examine the association between VPT birth and PLEs, separate models were fitted for each global and domain-specific PLE outcome, including total score, distressing items score, and distress severity score. In BIPP, multiple linear regression models were fitted with birth status as the main independent variable, adjusting for age, sex, and area-level socioeconomic deprivation. In ABCD, linear mixed-effects models were fitted with the same fixed effects and with random intercepts for recruitment site and family identifier, thereby accounting for the clustering of participants within sites and the inclusion of siblings or twins from the same family, consistent with recommended analytical practice for ABCD data^33^. Seven secondary models were fitted, all retaining the covariates from the primary model. Four models additionally included, separately, cognitive ability, total psychopathology, internalising symptoms, or externalising symptoms. Three further models included cognitive ability together with, respectively, total psychopathology, internalising symptoms, or externalising symptoms.

In an additional set of analyses, the BIPP and ABCD datasets were pooled in order to examine whether cohort membership was associated with PLEs independently of age and sex. Linear regression models were fitted in the combined sample with PQ-BC scores as the dependent variables and cohort as the main independent variable, first adjusting for age and sex, and then additionally for birth status. Finally, a cohort-by-birth status interaction term was included to assess whether the association between VPT birth and psychosis risk symptoms differed between cohorts.

To facilitate interpretation of effect sizes, standardised regression coefficients (standardised β) were additionally reported for the main regression models.

For the BIPP and pooled linear regression analyses, heteroscedasticity-consistent standard errors were estimated using the HC3 estimator, and statistical inference was based on robust Wald tests. ABCD mixed-effects models were estimated using maximum likelihood, with statistical significance of fixed effects evaluated using Satterthwaite-approximated degrees of freedom. Effect estimates are reported with standard errors, 95% confidence intervals, and p values. A two-sided *p* value < 0.05 was considered statistically significant for all analyses.

## 3. Results

### 3.1 Sample characteristics

Sample characteristics from BIPP and ABCD cohorts are shown in Table 1 and eResults 1, respectively. A descriptive comparison of characteristics available across both cohorts is provided in Table 2. In the BIPP cohort, children born VPT were older than term-born controls at assessment (*p* < 0.01). The groups did not differ significantly in sex or ethnicity. Caregivers of VPT children had lower educational attainment overall (*p* < 0.01), and the VPT group showed higher SDQ total and internalising scores than controls (both *p* < 0.01), whereas externalising scores did not differ significantly. VPT children also had lower age-adjusted full-scale IQ scores than term-born controls (*p* < 0.01).

**Table 1.** Demographic and clinical information of very preterm and control group in BIPP.

|  | **Control (N=72)** | **Very preterm (N=197)** | ***p* value** |
| --- | --- | --- | --- |
| **Age, y** | 9.22 ± 1.21 | 10.96 ± 1.71 | <0.01* |
| **Sex, n (%)** |  |  |  |
| Female | 42 (58.33%) | 106 (53.81%) | 0.60 |
| Male | 30 (41.67%) | 91 (46.19%) |  |
| **Gestational age, weeks** | 40.31 ± 1.19 | 29.86 ± 2.20 | <0.01* |
| **Birth weight, grams** | N.a. | 1306.10 ± 397.72 | N.a. |
| **Ethnicity, n (%)** |  |  |  |
| White / Caucasian | 38 (65.52%) | 82 (57.75%) | 0.10 |
| Black | 6 (10.34%) | 12 (8.45%) |  |
| Asian | 4 (6.90%) | 19 (13.38%) |  |
| Other, including Hispanic | 10 (17.24%) | 29 (20.42%) |  |
| **IMD income-deprivation decile** | 6.63 ± 2.98 | 5.97 ± 2.75 | <0.01* |
| **Highest education of caregiver, n (%)** |  |  |  |
| GCSEs | 2 (2.99%) | 17 (10.76%) | <0.01* |
| A Levels | 3 (4.48%) | 16 (10.13%) |  |
| Degree | 20 (29.85%) | 57 (36.08%) |  |
| Professional qualification | 13 (19.40%) | 29 (18.35%) |  |
| Master's degree or higher | 29 (43.28%) | 39 (24.68%) |  |
| **SDQ** |  |  |  |
| Total score | 7.31 ± 4.89 | 9.80 ± 6.38 | <0.01* |
| Internalising score | 2.85 ± 2.51 | 4.37 ± 3.65 | <0.01* |
| Externalising score | 4.46 ± 3.20 | 5.43 ± 3.84 | 0.14 |
| **IQ (Age-adjusted WISC, full scale)** | 108.80 ± 13.14 | 97.03 ± 15.99 | <0.01* |

Values are mean (SD) unless otherwise indicated. GCSE General Certificate of Secondary Education; IQ Intelligence Quotient; SDQ Strengths & Difficulties Questionnaire; WISC Wechsler Intelligence Scale for Children.

**Table 2.**
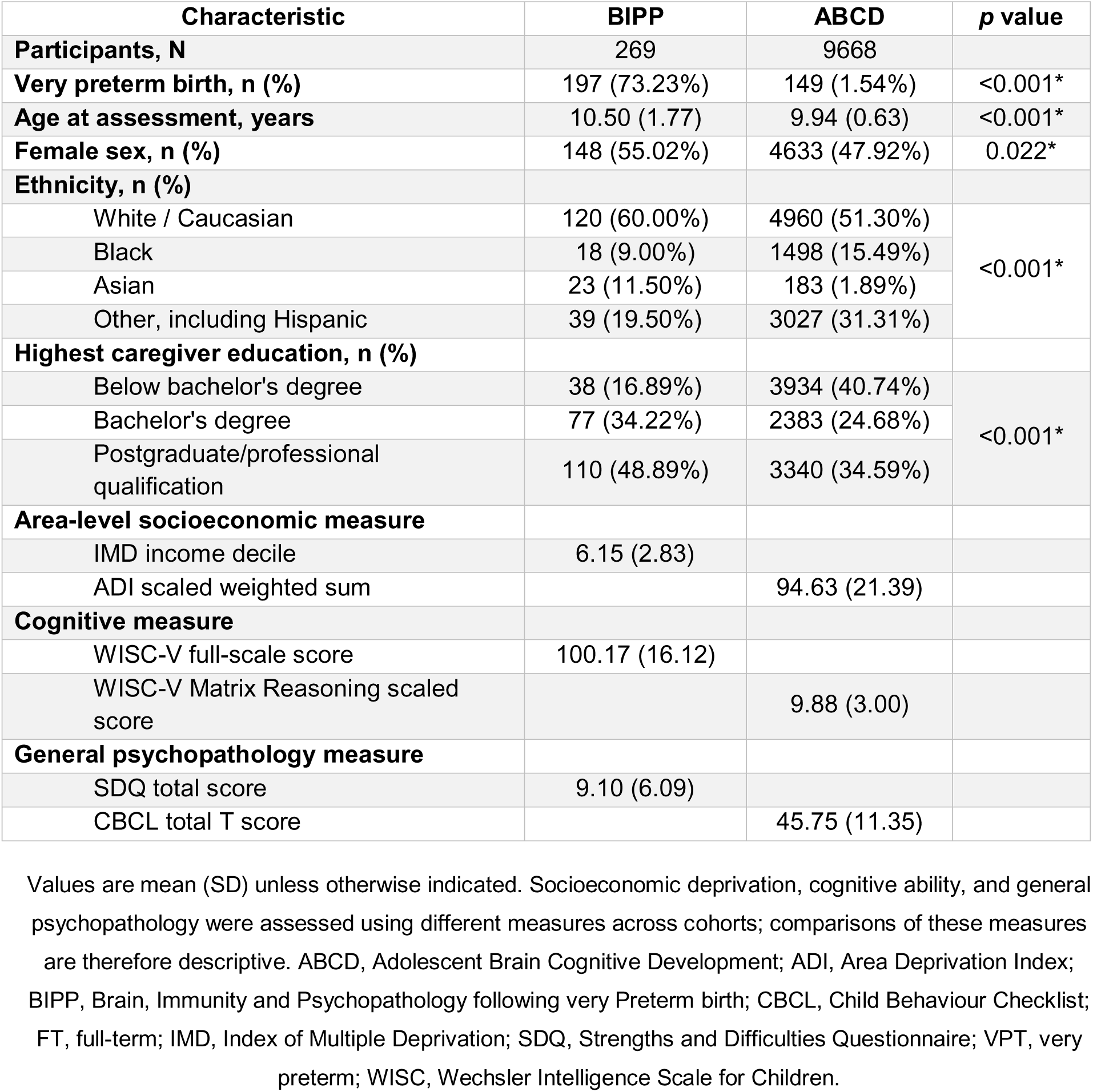
Descriptive comparison of demographic and clinical characteristics of the BIPP and ABCD cohorts.

| Characteristic | BIPP | ABCD | *p* value |
| --- | --- | --- | --- |
| **Participants, N** | 269 | 9668 |  |
| **Very preterm birth, n (%)** | 197 (73.23%) | 149 (1.54%) | <0.001* |
| **Age at assessment, years** | 10.50 (1.77) | 9.94 (0.63) | <0.001* |
| **Female sex, n (%)** | 148 (55.02%) | 4633 (47.92%) | 0.022* |
| **Ethnicity, n (%)** |  |  |  |
| White / Caucasian | 120 (60.00%) | 4960 (51.30%) | <0.001* |
| Black | 18 (9.00%) | 1498 (15.49%) |  |
| Asian | 23 (11.50%) | 183 (1.89%) |  |
| Other, including Hispanic | 39 (19.50%) | 3027 (31.31%) |  |
| **Highest caregiver education, n (%)** |  |  |  |
| Below bachelor's degree | 38 (16.89%) | 3934 (40.74%) | <0.001* |
| Bachelor's degree | 77 (34.22%) | 2383 (24.68%) |  |
| Postgraduate/professional qualification | 110 (48.89%) | 3340 (34.59%) |  |
| **Area-level socioeconomic measure** |  |  |  |
| IMD income decile | 6.15 (2.83) |  |  |
| ADI scaled weighted sum |  | 94.63 (21.39) |  |
| **Cognitive measure** |  |  |  |
| WISC-V full-scale score | 100.17 (16.12) |  |  |
| WISC-V Matrix Reasoning scaled score |  | 9.88 (3.00) |  |
| **General psychopathology measure** |  |  |  |
| SDQ total score | 9.10 (6.09) |  |  |
| CBCL total T score |  | 45.75 (11.35) |  |

Values are mean (SD) unless otherwise indicated. Socioeconomic deprivation, cognitive ability, and general psychopathology were assessed using different measures across cohorts; comparisons of these measures are therefore descriptive. ABCD, Adolescent Brain Cognitive Development; ADI, Area Deprivation Index; BIPP, Brain, Immunity and Psychopathology following very Preterm birth; CBCL, Child Behaviour Checklist; FT, full-term; IMD, Index of Multiple Deprivation; SDQ, Strengths and Difficulties Questionnaire; VPT, very preterm; WISC, Wechsler Intelligence Scale for Children.

In the ABCD cohort, children born VPT were slightly older than full-term controls at assessment (*p* = 0.03). The groups did not differ significantly in sex or ethnicity. As expected, children born VPT had lower birth weight than full-term controls (*p* < 0.01). The VPT group was more likely to be from lower-income households (*p* < 0.01), and to have caregivers with lower educational attainment; *p* <0.01. In contrast to the BIPP cohort, CBCL total, internalising, and externalising *t* scores did not differ significantly between groups. However, children born VPT had lower cognitive performance on the WISC-V Matrix Reasoning scaled scores than full-term controls (*p* < 0.01).

### 3.2 Associations between very preterm birth and PLEs in the BIPP cohort

In the primary BIPP analyses adjusted for age, sex, and socio-economic status, VPT birth was associated with a higher burden of PLEs in childhood compared with FT birth. Specifically, children born VPT had significantly higher PQ-BC total scores than term-born controls (β = 1.61, 95% CI 0.51 to 2.70, *p* = 0.004), as well as higher distressing items scores (β = 0.78, 95% CI 0.04 to 1.51, *p* = 0.039). The association with distress severity was in the same direction but did not reach conventional statistical significance (β = 2.31, 95% CI -0.04 to 4.65, *p* = 0.055).

When symptom domains were examined separately, the association between VPT birth and PLEs appeared to be primarily driven by perceptual abnormalities. Children born VPT showed higher perceptual abnormalities total scores (β = 0.82, 95% CI 0.35 to 1.29, *p* < 0.001), higher perceptual abnormalities distressing items scores (β = 0.49, 95% CI 0.18 to 0.81, *p* = 0.003), and higher distress severity related to perceptual abnormalities (β = 1.41, 95% CI 0.51 to 2.32, *p* = 0.002) after controlling for age, sex, and socio-economic status. By contrast, associations with unusual thought content were more limited. VPT birth was associated with higher unusual thought content total score (β = 0.60, 95% CI 0.05 to 1.16, *p* = 0.035), but not with the corresponding distressing items score or distress severity score (*p* > 0.05). No significant associations were observed between VPT birth and disorganised speech, whether indexed for total score, distressing items score, or distress severity score.

Across models, older age and higher socio-economic status were generally associated with fewer and less distressing PLEs. Higher IMD income-deprivation decile was consistently associated with lower scores across most global and domain-specific outcomes, whereas older age was more selectively associated with lower scores on total PLEs, unusual thought content, and perceptual abnormalities. Sex was not significantly associated with any of the PQ-BC outcomes. Detailed results for all regression models are provided in Table 3.

**Table 3.**
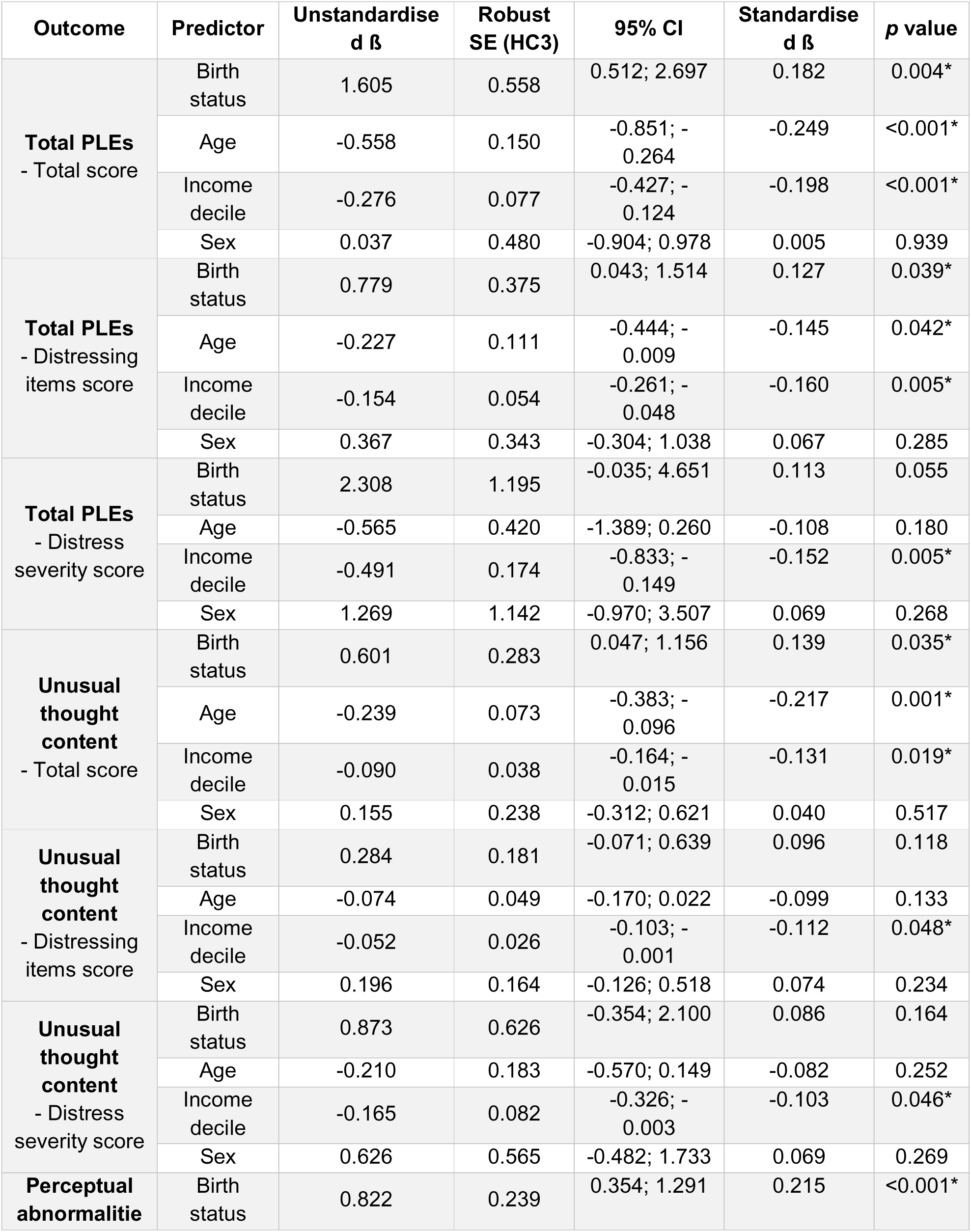
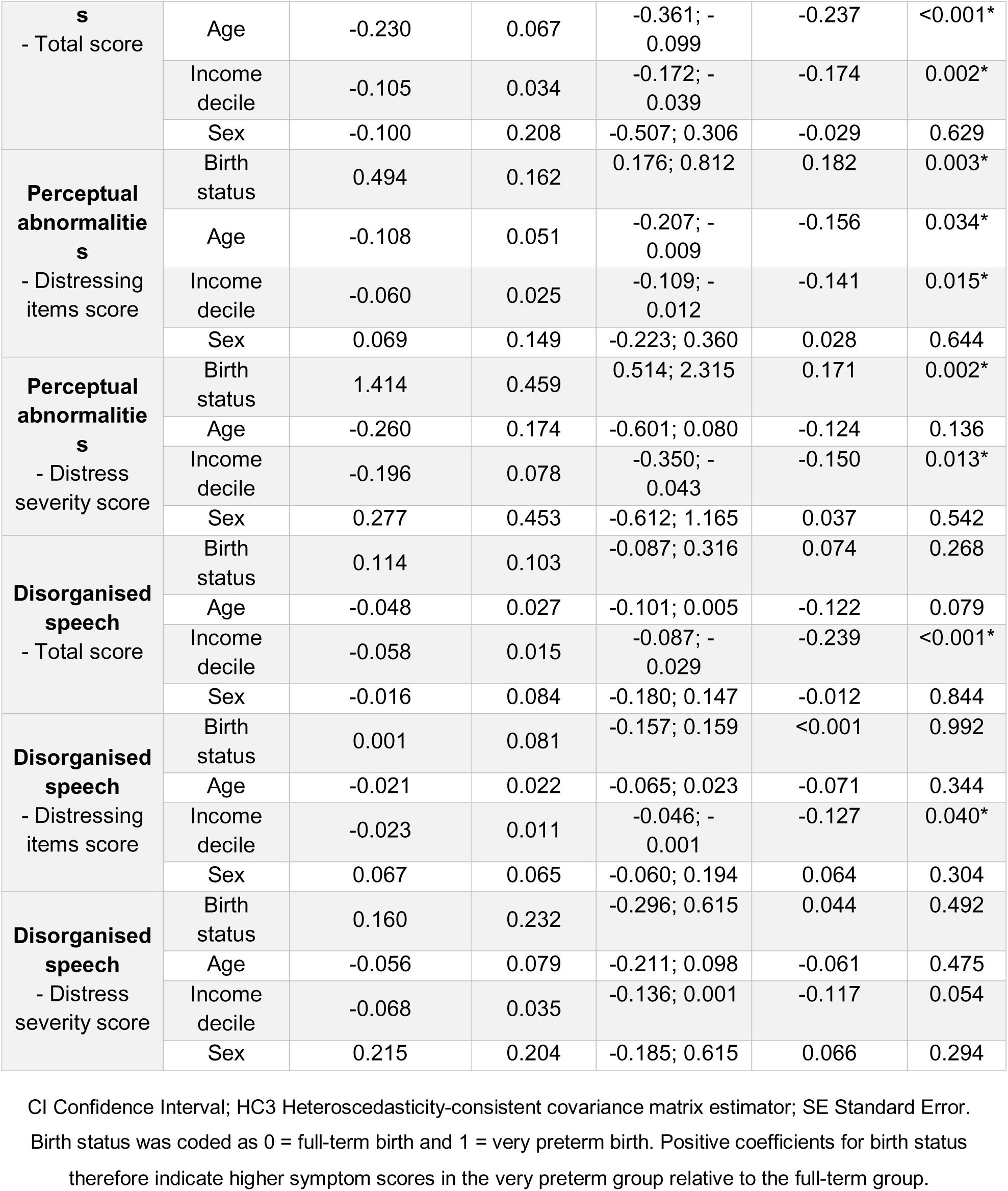
Adjusted regression models of psychotic-like experiences in childhood in the BIPP cohort.

### 3.3 Sensitivity analyses adjusting for cognitive ability and SDQ dimensions in the BIPP cohort

To assess the robustness of the primary findings, secondary models were fitted with additional adjustment for cognition and for general psychopathology, indexed separately by SDQ total difficulties, internalising difficulties, and externalising difficulties. Overall, the pattern of results remained similar to that observed in the primary analyses, although some associations were attenuated.

After adjustment for cognition, the association between VPT birth and overall PLEs burden remained significant for PQ-BC total score (β = 1.14, 95% CI 0.02 to 2.27, *p* = 0.048), but no longer for the global distress indices (*p* > 0.05). At the domain level, the most consistent associations continued to be observed for perceptual abnormalities, with VPT birth remaining associated with higher perceptual abnormalities total score (β = 0.67, 95% CI 0.19 to 1.15, *p* = < 0.001), distressing items score (β = 0.43, 95% CI 0.09 to 0.77, *p* = 0.015), and distress severity score (β = 1.19, 95% CI 0.24 to 2.13, *p* = 0.015). By contrast, associations with unusual thought content were no longer statistically significant after accounting for cognition, and no associations were observed for disorganised speech (all *p* > 0.05).

A similar pattern was observed in models additionally adjusting for general psychopathology. The association between VPT birth and PQ-BC total score remained significant when the models included SDQ total difficulties (β = 1.20, 95% CI 0.01 to 2.39, *p* = 0.049), internalising difficulties (β = 1.24, 95% CI 0.07 to 2.42, *p* = 0.040), or externalising difficulties (β = 1.37, 95% CI 0.21 to 2.53, *p* = 0.022), whereas associations with the global distress outcomes were attenuated. Across all three SDQ-adjusted models, the association between VPT birth and perceptual abnormalities remained significant for total score and distress score and was also retained for distress severity in the models including internalising and externalising difficulties, while showing borderline significance in the model including total difficulties. In contrast, associations with unusual thought content were attenuated and no longer statistically significant, and there was again no evidence of an association with disorganised speech (all *p* > 0.05).

When cognitive ability and SDQ total difficulties were included simultaneously, the association between VPT birth and overall PQ-BC total score was attenuated and no longer statistically significant, and the same was true for the global distress indices (all *p* > 0.05). At the domain level, VPT birth remained associated only with perceptual abnormalities total score (β = 0.55, 95% CI 0.04 to 1.07, *p* = 0.036), whereas associations with perceptual distressing items and perceptual distress severity were reduced to borderline significance. Similarly, when cognition and internalising difficulties were entered simultaneously, associations with global PQ-BC outcomes were no longer statistically significant, while the association with perceptual abnormalities total score remained significant and associations with perceptual distressing items and perceptual distress severity were attenuated to borderline significance. By contrast, in models including both cognition and externalising difficulties, the association with overall PQ-BC total score was attenuated to borderline significance, whereas the association with perceptual abnormalities remained significant across all three indexes.

Detailed results of the models additionally adjusted for cognitive ability, SDQ total difficulties, internalising difficulties, and externalising difficulties are provided in eResults 2-8. The pattern of findings across primary and secondary models is summarised in Figure 1.

**Figure 1.**
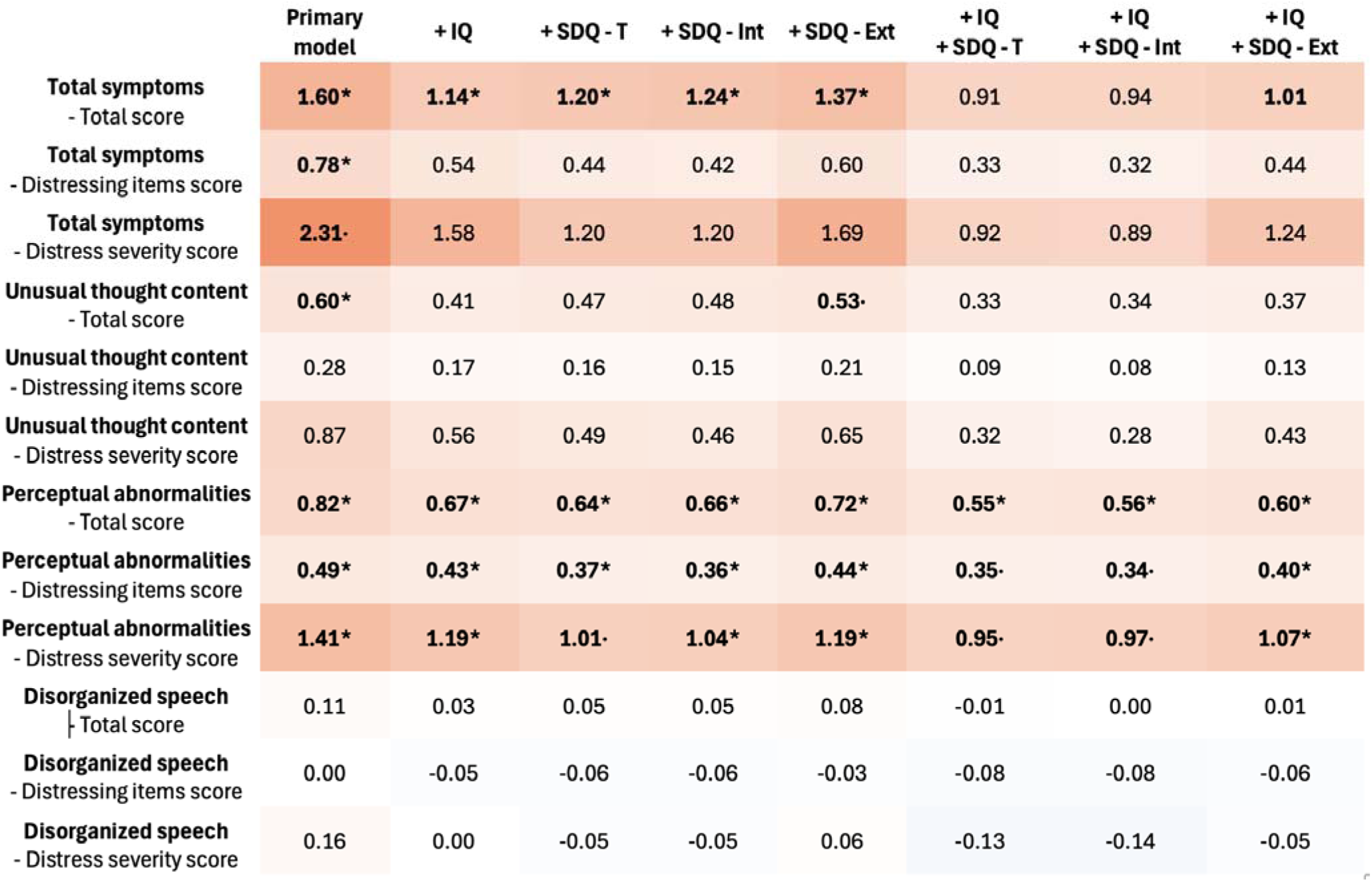
Heat map summarising the association between very preterm birth and psychotic-like experiences across primary and secondary models in the BIPP cohort. Cells represent regression coefficients (β) for birth status from linear regression models examining global and domain-specific PQ-BC outcomes. The primary model was adjusted for age, sex, and socio-economic status. Secondary models included additional adjustment for IQ, SDQ total difficulties, SDQ internalising difficulties, SDQ externalising difficulties, and combined models including IQ together with each SDQ measure. Warmer colors indicate stronger positive associations with very preterm birth, cooler colors indicate weaker or negative associations, and white indicates coefficients close to zero. Asterisks indicate statistically significant associations (* *p* < 0.05), and dots indicate trend-level associations (· *p* < 0.10).

### 3.4 Validation analyses in the ABCD cohort

In the primary ABCD analyses adjusted for age, sex, and socio-economic status, VPT birth was not significantly associated with overall PLEs in childhood. No significant association was observed for PQ-BC total score (β = -0.26, 95% CI -0.84 to 0.32, *p* = 0.384), distressing items score (β = 0.046, 95% CI -0.33 to 0.42, *p* = 0.808) and distress severity (β = 0.25, 95% CI -0.96 to 1.46, *p* = 0.686). When symptom domains were examined separately, no significant associations were observed between VPT birth and unusual thought content, perceptual abnormalities, or disorganised speech across total, distressing items, or distress severity scores. Thus, unlike in the BIPP cohort, the domain-specific pattern observed in the primary analyses were not clearly replicated in ABCD.

Across models, older age and lower neighbourhood deprivation, indicated by lower ADI scores, were generally associated with fewer PLEs, whereas female sex was associated with lower scores across several global and domain-specific outcomes. Further adjustment for cognitive ability, general psychopathology, and their combination did not materially alter the overall pattern of findings: associations between VPT birth and global or domain-specific PLEs remained non-significant across all secondary models. By contrast, lower cognitive ability and higher levels of general psychopathology were consistently associated with greater PLEs burden across global and domain-specific PQ-BC outcomes. Detailed results for the primary and secondary ABCD models are provided in Table 4 and eResults 9-15.

**Table 4.**
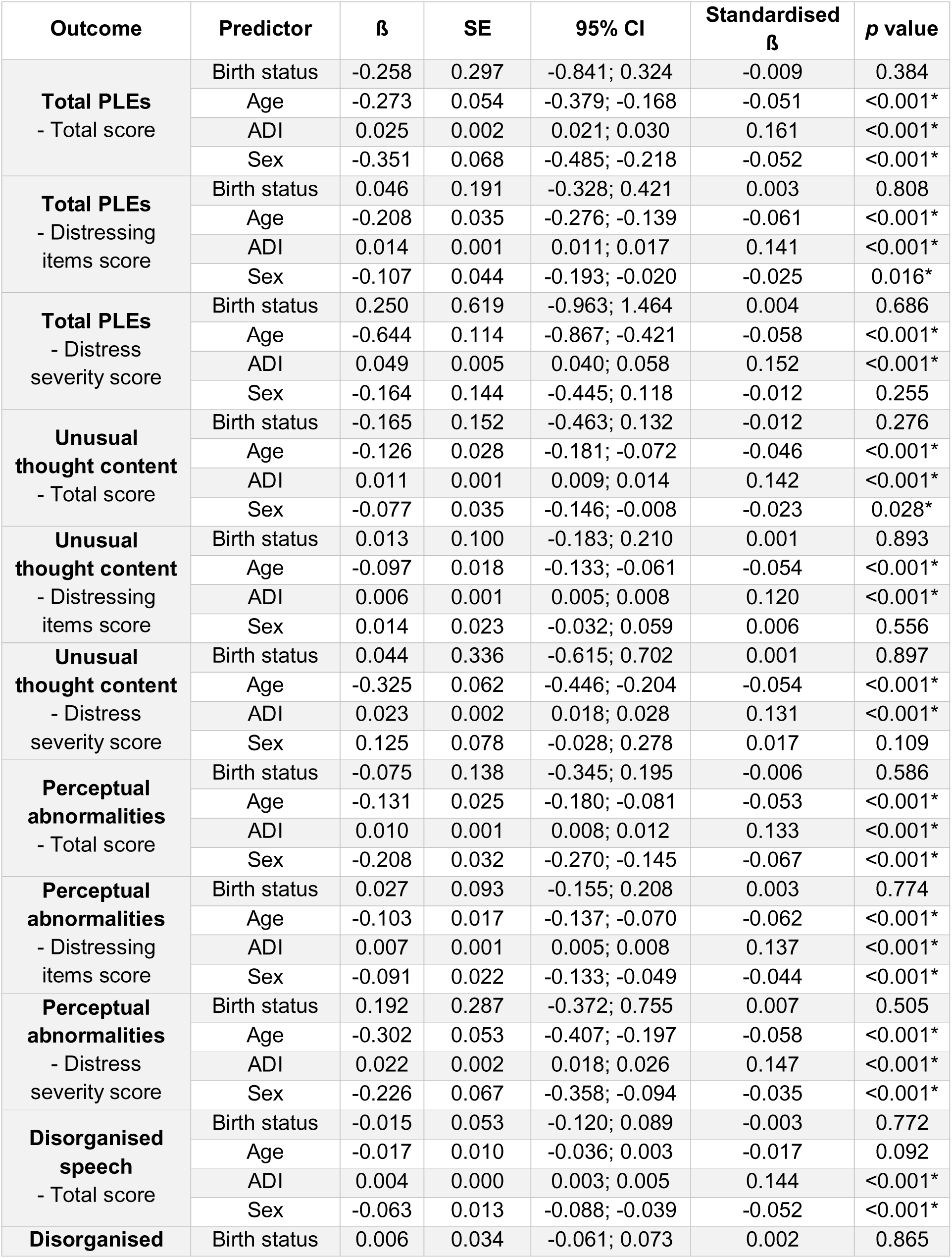
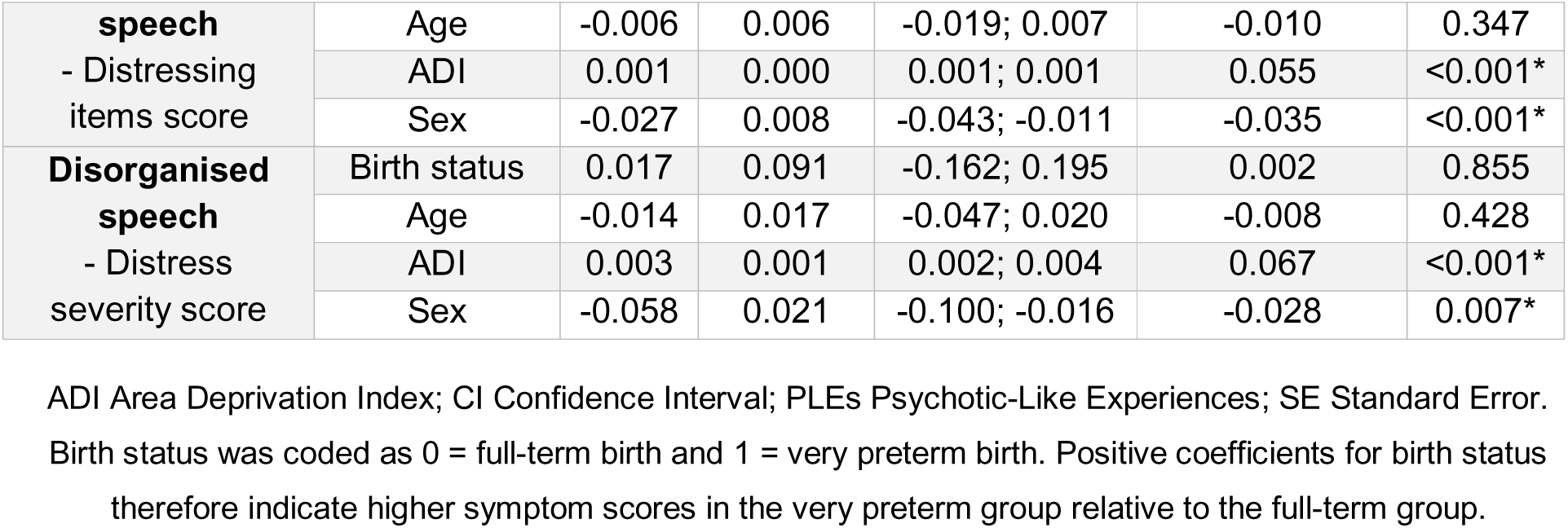
Adjusted regression models of psychotic-like experiences in childhood in the ABCD cohort.

### 3.5 Pooled analyses across the BIPP and ABCD cohorts

In pooled analyses combining the BIPP and ABCD cohorts, cohort membership was significantly associated with total PLEs after adjustment for age and sex. Relative to BIPP, membership in the ABCD cohort was associated with lower total PQ-BC scores (β = -1.25, 95% CI - 1.73 to -0.78, *p* < 0.001), lower distressing items scores (β = -0.72, 95% CI -1.05 to -0.39, *p* < 0.001), and lower distress severity scores (β = -2.32, 95% CI -3.41 to -1.22, *p* < 0.001). A similar pattern was observed across most domain-specific outcomes, with ABCD participants generally showing lower scores than BIPP participants for unusual thought content, perceptual abnormalities, and disorganised speech. Across outcomes, older age was consistently associated with fewer PLEs (see eResults 16).

When birth status was added to the pooled models, the association between cohort membership and PLEs was attenuated but remained significant for several global and domain-specific outcomes. Preterm birth was independently associated with higher total PLEs scores (β = 0.55, 95% CI 0.10 to 0.99, *p* = 0.018), distressing items scores (β = 0.43, 95% CI 0.13 to 0.73, *p* = 0.006), and distress severity scores (β = 1.40, 95% CI 0.41 to 2.39, *p* = 0.006), as well as with higher unusual thought content distressing items and distress severity, and higher perceptual abnormalities across total, distressing items, and distress severity scores. By contrast, no significant associations with preterm birth were observed for disorganised speech outcomes. These findings suggest that differences between cohorts were only partly explained by differences in birth status composition (see eResults 17).

In models including a cohort-by-birth status interaction, preterm birth was associated with higher total PLEs scores within the reference cohort, BIPP, including total score, distressing items score, and distress severity score. Significant positive associations with preterm birth were also observed for perceptual abnormalities across all three indices, whereas associations with unusual thought content were weaker and limited to trend-level effects for total and distress-related outcomes, and no significant associations were observed for disorganised speech. Importantly, the interaction term indicated that the association between preterm birth and PLEs was weaker in ABCD than in BIPP for perceptual abnormalities total score, distressing items score, and distress severity score, while the interaction for total PLEs score was at trend level. No significant interaction effects were observed for unusual thought content or disorganised speech. Overall, these pooled analyses suggest that the association between VPT birth and PLEs observed in BIPP, particularly for perceptual abnormalities, was attenuated in ABCD (see Table 5).

**Table 5.**
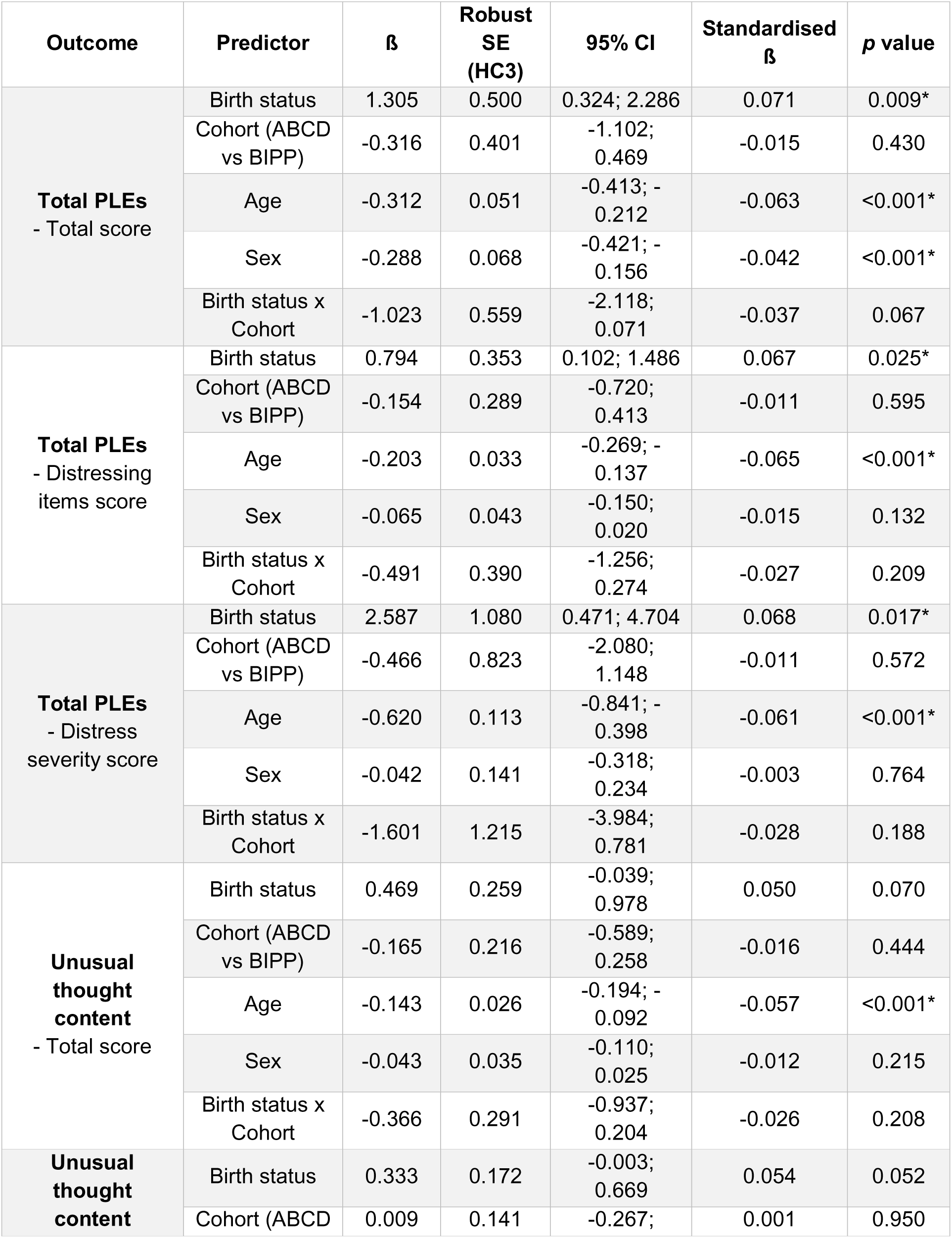
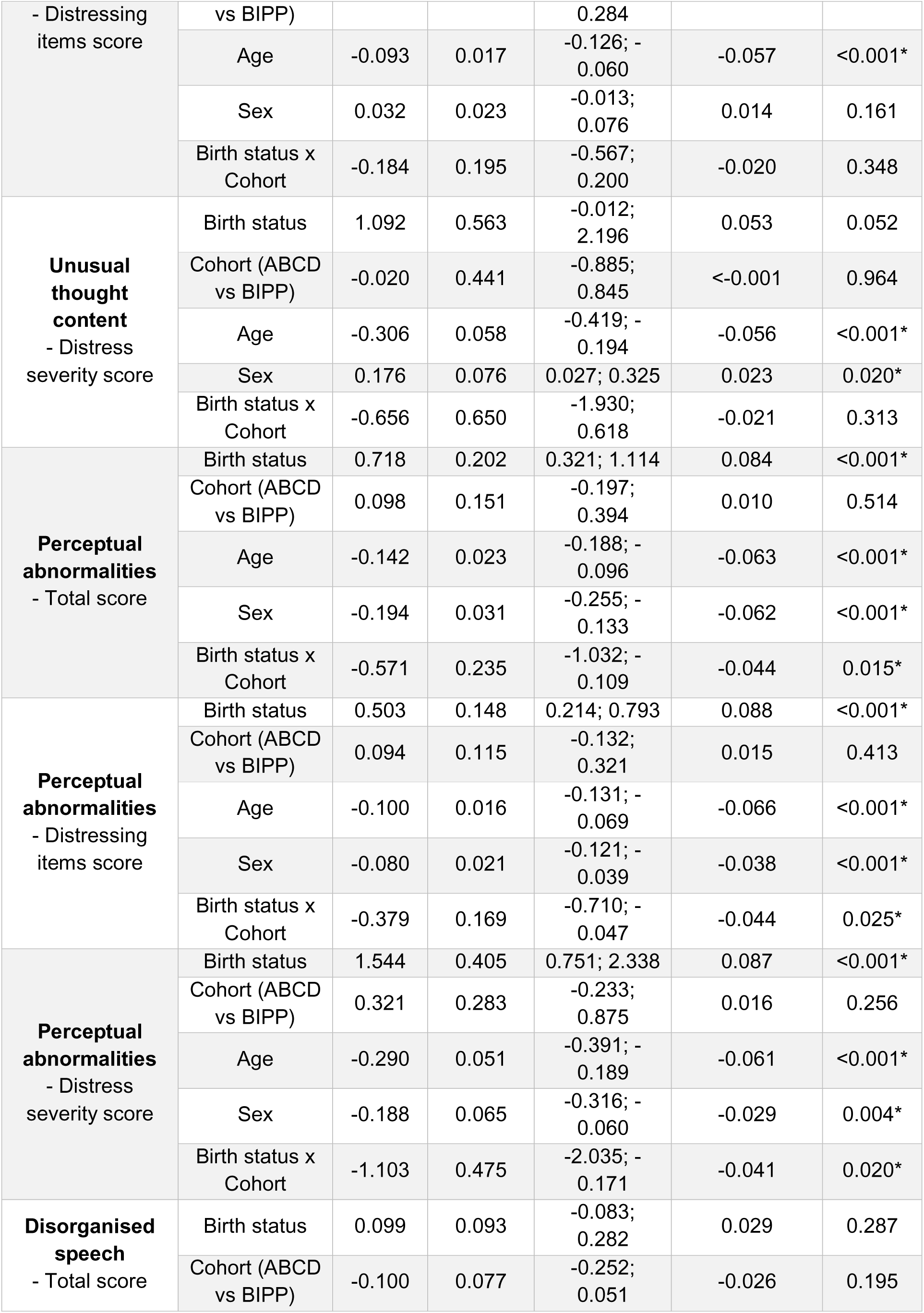
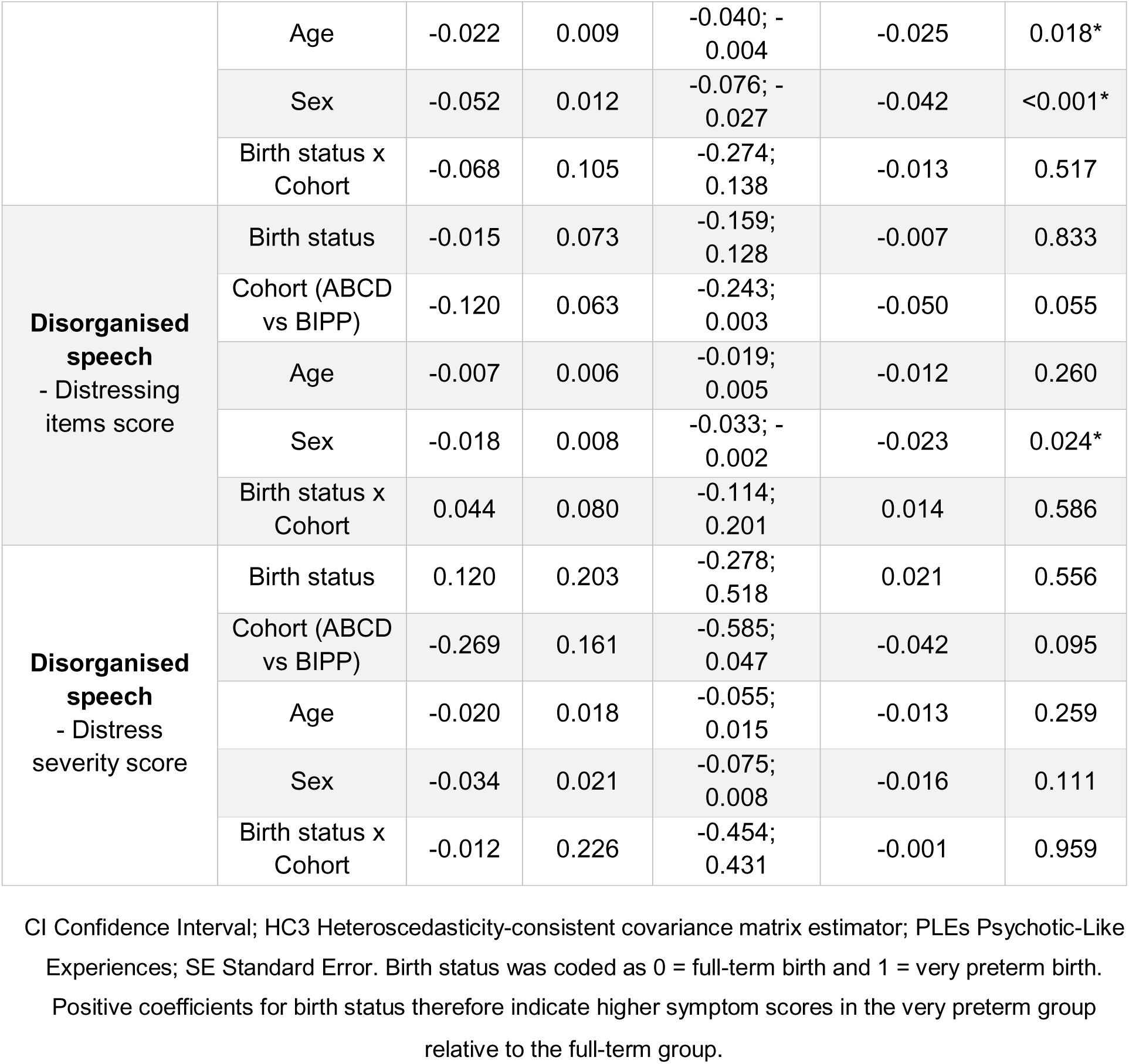
Pooled analyses across the BIPP and ABCD cohorts. Model including cohort-by- birth status interaction, adjusted for age and sex.

## 4. Discussion

In this study, children born very preterm recruited from a clinical birth cohort (BIPP) showed a greater burden of psychotic-like experiences in childhood, particularly perceptual abnormalities, although these findings were not replicated in a population-based cohort (ABCD). To our knowledge, this is the first study to investigate psychotic-like experiences in children born very preterm.

Several important findings emerged. First, the association observed in BIPP suggests increased early vulnerability to psychosis-spectrum outcomes among children born VPT. Importantly, this association was not fully accounted for by differences in cognitive ability or broader psychopathology. This is particularly relevant given that cognitive deficits are a well-established feature of psychotic disorders^34^, including their early stages^14,15^, and are also consistently reported in individuals born VPT^35^. Likewise, general psychopathology is closely linked to psychosis-spectrum phenomena at the population level^10^ and is more common among children born VPT^2,36,37^. The fact that the association between VPT and PLEs persisted after accounting for these factors suggests that it may reflect a partially distinct dimension of vulnerability.

Another noteworthy finding was that the association between VPT birth and PLEs was not evenly distributed across symptom domains but appeared to be more specifically driven by perceptual abnormalities rather than unusual thought content or disorganised speech. This pattern remained consistent across models and again persisted after adjustment for cognitive ability and broader psychopathology. The identification of such a domain-specific profile is particularly informative, as PLEs are not a unitary construct and different dimensions may partly reflect distinct mechanisms of risk^9^. In the context of VPT birth, the prominence of perceptual abnormalities may indicate a pathway more closely related to neurodevelopmental alterations in perceptual processing or sensory integration^38,39^. More broadly, these findings support the notion that VPT birth may be associated with specific psychopathological phenotypes, rather than with uniform elevation in risk across all symptom domains. This is consistent with evidence in other neurodevelopmental conditions, where preterm-born individuals show a more pronounced liability for inattentive, but not hyperactive-impulsive, ADHD symptoms^40^, as well as a more specific autism-spectrum profile involving circumscribed social deficits but fewer restricted and repetitive behaviours^41^.

We were unable to replicate these results in the independent population-based ABCD cohort, where VPT birth was not significantly associated with overall or domain-specific PLEs. We tentatively suggest that the lack of replication may, at least in part, relate to the composition of the ABCD cohort, and particularly to the characteristics of its VPT subsample. First, ABCD excluded children born extremely preterm (<28 weeks’ gestation)^21^, who are likely to represent the subgroup at highest neurodevelopmental and psychiatric risk^42^. Second, because recruitment took place in childhood rather than at birth, as in BIPP, the study may have been less likely to capture those children born VPT with more marked functional difficulties, who may have faced greater barriers to participation in a community-based study^43^. Taken together, these features may have resulted in a particularly resilient or better-functioning VPT subgroup, thereby reducing the likelihood of replicating the findings observed in BIPP.

In fact, several observations suggest that the ABCD VPT subgroup may not be fully representative of the wider VPT population. In contrast to both the BIPP cohort^2^ and the wider literature^44^, children born VPT in ABCD did not show elevated levels of overall, internalising, or externalising psychopathology relative to FT controls. This absence of elevated psychopathology is notable, as children born VPT are generally reported to show increased rates of emotional, behavioural, and broader psychiatric difficulties compared with their FT peers^2,37,45^. Taken together, these findings suggest that the ABCD VPT subgroup may represent a relatively normative or resilient segment of the VPT population, consistent with suggestions from previous ABCD-based studies in other research contexts^46^.

A range of mechanisms may underlie the association between VPT birth and PLEs, which is unlikely to be attributable to a single pathway. A first possibility is that this association reflects atypical brain development following VPT birth. Being born preterm exposes the immature brain to the extrauterine environment at a highly sensitive stage, when key neurodevelopmental processes are still unfolding^47^, and this may have enduring consequences for later brain and behavioural functioning. Indeed, VPT birth has been linked not only to severe neonatal brain injury^48^, but also to more subtle and widespread alterations in brain structure and organisation, including alterations affecting white matter, cortical development, and large-scale neural networks^49–51^, many of which overlap with neural systems implicated in psychosis^6^. Within this framework, the pattern observed in the present study – namely a greater burden of PLEs in the BIPP cohort, with the clearest signal emerging for perceptual abnormalities – may point to a pathway related to altered perceptual processing or sensory integration, which has already been described in populations born preterm^52,53^.

Neonatal adversity may also contribute to later psychosis-spectrum vulnerability through both direct and indirect pathways. Directly, complications associated with VPT birth, such as hypoxic–ischemic events^54^, inflammation^55^, or repeated exposure to pain and intensive care^56^, may disrupt early brain maturation and the development of neural systems relevant to salience processing, sensory integration, and cognitive control^57^. Indirectly, these early adversities may contribute to later attentional^58^, social^59^, motor^60^, or sensory-integration difficulties, which could in turn increase vulnerability to subclinical psychotic experiences. In fact, the association between VPT birth and prodromal symptoms was attenuated after adjustment for IQ and broader psychopathology, suggesting that these factors contribute to, but do not fully account for, the observed relationship. Taken together, these findings are consistent with the broader view that VPT birth confers a general neurodevelopmental vulnerability associated with multiple psychiatric outcomes, of which psychosis-spectrum symptoms may be one expression.

Studying these phenomena in childhood is particularly important, as it offers a window into very early developmental trajectories of vulnerability. In the context of VPT birth, this may help clarify whether psychosis-spectrum risk is already detectable before adolescence, when more overt clinical manifestations typically begin to emerge^61^, and may improve our understanding of the medium- and long-term psychiatric prognosis of this population. These findings also have important clinical implications, highlighting the need for sustained attention to the mental health of children born VPT beyond the neurodevelopmental difficulties that are more typically recognised^36,62^. Although psychosis-spectrum experiences should not be interpreted as indicating an inevitable trajectory towards psychotic disorder, they may represent an additional dimension of vulnerability that warrants careful long-term monitoring, particularly when subtle perceptual or unusual experiences are persistent, distressing, or accompanied by broader emotional, behavioural, or cognitive difficulties. From a research perspective, longitudinal studies will be essential to determine the clinical significance of these early manifestations, including whether they persist over time, predict later psychopathology, or instead reflect transient developmental variation within this population^11^.

This study has several limitations. First, PLEs were assessed using a self-report questionnaire rather than a clinician-administered assessment. Self-report measures in childhood are subject to inherent limitations, including acquiescence, difficulties in item comprehension, and developmental variation in the meaning of unusual experiences^12^. Nonetheless, the PQ-BC is a well-validated and widely used measure of psychotic-like experiences in children and provides a useful dimensional index of early psychosis-spectrum symptoms^28^. Second, the comparison between the BIPP and ABCD cohorts was not exact, as certain measures were not directly comparable across studies. General psychopathology and cognitive ability were assessed using different instruments across cohorts (SDQ vs CBCL; WISC full-scale score vs WISC-V Matrix Reasoning scaled score). Although these measures assess related domains, they are not directly equivalent, which may have introduced measurement heterogeneity. Third, in ABCD, VPT birth was defined based on parent-report rather than clinically recorded gestational age. Although maternal recall is generally considered reasonably reliable for perinatal variables of this kind, this approach remains less precise than direct obstetric data^63^. Finally, the cross-sectional design precludes conclusions regarding the persistence, developmental course, and long-term clinical significance of these early experiences.

In conclusion, our findings suggest that VPT birth is associated with increased vulnerability to psychotic-like experiences in childhood, particularly in the domain of perceptual abnormalities. The lack of clear replication in the ABCD cohort may reflect differences in the composition of its very preterm subgroup, which may not be fully representative of the broader VPT population recruited from clinical settings. Further research is needed to better define this potentially specific phenotype and to establish the developmental and clinical significance of these early symptoms over time.

## Data availability statement

Data from the BIPP Study that support the findings of this manuscript are available on request from the corresponding author. The data are not publicly available due to ethical restrictions. Data from the ABCD Study (https://abcdstudy.org) is publicly available and can be accessed via the new NIH Brain Development Cohorts (NBDC) Data Hub.

## Acknowledgements

The authors would like to sincerely thank all the participants and their families. This work would not have been possible without their collective contributions.

## Ethical standard

The ePrime study was approved by the Hammersmith and Queen Charlotte’s Research Ethics Committee (REC: 09/H0707/98), BIPP by the South East Research Ethics Committee (REC: 19/LO/1940), and the Stanmore Research Ethics Committee (REC: 18/LO/0048). Written informed assent and consent to participate in this study was provided by the participants and their legal guardian/next of kin prior to data collection.

