## Supplementary Material for "Psychotic-like experiences in children born very preterm: evidence from clinical and population-based cohorts"

**Prodromal symptoms in children born very preterm: Findings from Brain, Immunity and Psychopathology following very Preterm birth (BIPP) and Adolescent Brain Cognitive Development (ABCD) Studies**

- SUPPLEMENTARY MATERIAL -

### **eMethods 1**. Classification of PQ-BC items into symptom domains based on the SIPS

**Unusual thought content domain**

Item 1. *Do familiar surroundings sometimes seem strange, confusing, threatening or unreal to you?*
Item 4. *Have you had experiences with telepathy, psychic forces, or fortune telling?*
Item 5. *Have you felt that you are not in control of your own ideas or thoughts?*
Item 7. *Do you have strong feelings or beliefs about being unusually gifted or talented in some way?*
Item 8. *Do you feel that other people are watching you or talking about you?*
Item 11. *Have you had the sense that some person or force is around you, although you couldn’t see anyone?*
Item 13. *Have you ever felt that you don't exist, the world does not exist, or that you are dead?*
Item 15. *Do you hold beliefs that other people would find unusual or bizarre?*
Item 18. *Do you find yourself feeling mistrustful or suspicious of other people?*

**Perceptual abnormalities domain**

Item 2. *Have you heard unusual sounds like banging, clicking, hissing, clapping or ringing in your ears?*
Item 3. *Do things that you see appear different from the way they usually do (brighter or duller, larger or smaller, or changed in some other way)?*
Item 9. *Do you sometimes get strange feelings on or just beneath your skin, like bugs crawling?*
Item 10. *Do you sometimes feel suddenly distracted by distant sounds that you are not normally aware of?*
Item 16. *Do you feel that parts of your body have changed in some way, or that parts of your body are working differently?*
Item 17. *Are your thoughts sometimes so strong that you can almost hear them?*
Item 19. *Have you seen unusual things like flashes, flames, blinding light, or geometric figures?*
Item 20. *Have you seen things that other people can't see or don't seem to see?*

**Disorganized speech domain**

Item 6. *Do you have difficulty getting your point across, because you ramble or go off the track a lot when you talk?*
Item 21. *Do people sometimes find it hard to understand what you are saying?*

### **eResults 1**. Demographic and clinical characteristics of ABCD sample

|  | FT (N=9519) | VPT (N=149) | *p* value |
| --- | --- | --- | --- |
| Age, y | 9.93 ± 0.63 | 10.04 ± 0.65 | 0.03* |
| Gender, n (%) |  |  |  |
| Female | 4561 (47.91%) | 72 (48.32%) | 0.99 |
| Male | 4958 (52.09%) | 77 (51.68%) |  |
| Gestational age, week | N.a. | 29.15 ± 1.63 | N.a. |
| Birth weight, kilograms | 3.17 ± 0.54 | 1.62 ± 0.67 | <0.01* |
| Race / Ethnicity, n (%) |  |  |  |
| White | 4884 (51.38%) | 70 (46.98%) | 0.25 |
| Black | 1466 (15.42%) | 30 (20.13%) |  |
| Hispanic | 2015 (21.20%) | 37 (24.83%) |  |
| Asian | 180 (1.89%) | 2 (1.34%) |  |
| Other | 961 (10.11%) | 10 (6.71%) |  |
| Area Deprivation Index | 94.53 ± 21.47 | 101.40 ± 14.02 | <0.01* |
| Highest education of caregiver, n (%) |  |  |  |
| < High school | 516 (5.43%) | 10 (6.71%) | <0.01* |
| High school diploma or GED | 921 (9.70%) | 28 (18.79%) |  |
| Some college or associate degree | 2415 (25.43%) | 37 (24.83%) |  |
| Bachelor’s degree | 2350 (24.75%) | 31 (20.81%) |  |
| Postgraduate degree | 3293 (34.68%) | 43 (28.86%) |  |
| CBCL |  |  |  |
| Total *t* score | 45.76 ± 11.35 | 45.13 ± 10.87 | 0.46 |
| Internalizing *t* score | 48.38 ± 10.61 | 48.52 ± 10.40 | 0.78 |
| Externalizing *t* score | 45.65 ± 10.30 | 44.30 ± 10.12 | 0.11 |
| Cognitive Ability (WISC-V Matrix Reasoning scaled score) | 9.89 ± 3.00 | 8.94 ± 2.67 | <0.01* |

ABCD Adolescent Brain Cognitive Development; CBCL Child Behaviour Checklist; FT Full Term; GED General Education Development; VPT Very Preterm

### **eResults 2**. Adjusted regression models of psychotic-like experiences in childhood in the BIPP cohort, additionally adjusted for Cognitive Ability

| Outcome | Predictor | ß | Robust SE (HC3) | 95% CI | *p* value |
| --- | --- | --- | --- | --- | --- |
| Total PLEs  - Total score | Birth status | 1.144 | 0.575 | 0.017; 2.271 | 0.048* |
|  | Age | -0.657 | 0.155 | -0.960; -0.354 | <0.001* |
|  | Income decile | -0.241 | 0.077 | -0.393; -0.089 | 0.002* |
|  | Sex | -0,140 | 0.492 | -1.103; 0.823 | 0.776 |
|  | Cognitive Ability | -0.049 | 0.016 | -0.081; -0.017 | 0.003* |
| Total PLEs  - Distressing items score | Birth status | 0.535 | 0.411 | -0.270; 1.340 | 0.194 |
|  | Age | -0.298 | 0.117 | -0.527; -0.069 | 0.011* |
|  | Income decile | -0.132 | 0.053 | -0.237; -0.027 | 0.014* |
|  | Sex | 0.249 | 0.360 | -0.457; 0.955 | 0.489 |
|  | Cognitive Ability | -0.031 | 0.012 | -0.056; -0.007 | 0.013* |
| Total PLEs  - Distress severity score | Birth status | 1.581 | 1.294 | -0.955; 4.117 | 0.223 |
|  | Age | -0.788 | 0.417 | -1.606; 0.030 | 0.060 |
|  | Income decile | -0.410 | 0.175 | -0.754; -0.067 | 0.020* |
|  | Sex | 0.935 | 1.185 | -1.388; 3.258 | 0.431 |
|  | Cognitive Ability | -0.101 | 0.038 | -0.175; -0.026 | 0.009* |
| Unusual thought content  - Total score | Birth status | 0.405 | 0.295 | -0.172; 0.983 | 0.170 |
|  | Age | -0.273 | 0.076 | -0.423; -0.123 | <0.001* |
|  | Income decile | -0.077 | 0.038 | -0.152; -0.002 | 0.046* |
|  | Sex | 0.087 | 0.246 | -0.396; 0.569 | 0.725 |
|  | Cognitive Ability | -0.018 | 0.008 | -0.034; -0.002 | 0.032* |
| Unusual thought content  - Distressing items score | Birth status | 0.166 | 0.199 | -0.224; 0.556 | 0.404 |
|  | Age | -0.104 | 0.052 | -0.206; -0.001 | 0.049* |
|  | Income decile | -0.044 | 0.026 | -0.094; 0.006 | 0.088 |
|  | Sex | 0.141 | 0.173 | -0.198; 0.479 | 0.416 |
|  | Cognitive Ability | -0.013 | 0.006 | -0.025; -0.001 | 0.035* |
| Unusual thought content  - Distress severity score | Birth status | 0.556 | 0.691 | -0.798; 1.909 | 0.422 |
|  | Age | -0.290 | 0.185 | -0.653; 0.073 | 0.119 |
|  | Income decile | -0.147 | 0.082 | -0.308; 0.015 | 0.076 |
|  | Sex | 0.465 | 0.589 | -0.690; 1.619 | 0.431 |
|  | Cognitive Ability | -0.034 | 0.029 | -0.074; 0.005 | 0.091 |
| Perceptual abnormalities  - Total score | Birth status | 0.667 | 0.245 | 0.186; 1.147 | 0.007* |
|  | Age | -0.259 | 0.068 | -0.393; -0.126 | <0.001* |
|  | Income decile | -0.098 | 0.035 | -0.166; -0.030 | 0.005* |
|  | Sex | -0.146 | 0.211 | -0.559; 0.267 | 0.488 |
|  | Cognitive Ability | -0.016 | 0.007 | -0.029; -0.003 | 0.016* |
| Perceptual abnormalities  - Distressing items score | Birth status | 0.428 | 0.174 | 0.086; 0.769 | 0.015* |
|  | Age | -0.130 | 0.053 | -0.234; -0.026 | 0.015* |
|  | Income decile | -0.055 | 0.025 | -0.103; -0.007 | 0.027* |
|  | Sex | 0.033 | 0.156 | -0.273; 0.338 | 0.833 |
|  | Cognitive Ability | -0.010 | 0.005 | -0.020; 0.000 | 0.059 |
| Perceptual abnormalities  - Distress severity score | Birth status | 1.186 | 0.483 | 0.240; 2.133 | 0.015* |
|  | Age | -0.343 | 0.173 | -0.682; -0.003 | 0.049* |
|  | Income decile | -0.169 | 0.078 | -0.322; -0.015 | 0.032* |
|  | Sex | 0.166 | 0.467 | -0.750; 1.082 | 0.723 |
|  | Cognitive Ability | -0.037 | 0.015 | -0.066; -0.009 | 0.011* |
| Disorganized speech  - Total score | Birth status | 0.031 | 0.104 | -0.173; 0.235 | 0.767 |
|  | Age | -0.072 | 0.028 | -0.127; -0.017 | 0.011 |
|  | Income decile | -0.049 | 0.015 | -0.078; -0.020 | 0.001 |
|  | Sex | -0.066 | 0.085 | -0.234; 0.101 | 0.438 |
|  | Cognitive Ability | -0.010 | 0.003 | -0.015; -0.004 | <0.001* |
| Disorganized speech  - Distressing items score | Birth status | -0.046 | 0.085 | -0.213; 0.121 | 0.591 |
|  | Age | -0.033 | 0.024 | -0.080; 0.014 | 0.170 |
|  | Income decile | -0.019 | 0.012 | -0.042; 0.004 | 0.113 |
|  | Sex | 0.046 | 0.068 | -0.087; 0.179 | 0.500 |
|  | Cognitive Ability | -0.005 | 0.002 | -0.009; 0.000 | 0.054 |
| Disorganized speech  - Distress severity score | Birth status | -0.003 | 0.240 | -0.473; 0.467 | 0.989 |
|  | Age | -0.098 | 0.084 | -0.264; 0.067 | 0.245 |
|  | Income decile | -0.048 | 0.038 | -0.123; 0.027 | 0.212 |
|  | Sex | 0.152 | 0.217 | -0.273; 0.576 | 0.485 |
|  | Cognitive Ability | -0.018 | 0.009 | -0.035; -0.001 | 0.040 |

CI Confidence Interval; HC3 Heteroscedasticity-consistent covariance matrix estimator; SE Standard Error. Birth status was coded as 0 = full-term birth and 1 = very preterm birth. Positive coefficients for birth status therefore indicate higher symptom scores in the very preterm group relative to the full-term group.

### **eResults 3**. Adjusted regression models of psychotic-like experiences in childhood in the BIPP cohort, additionally adjusted for SDQ – Total difficulties score

| Outcome | Predictor | ß | Robust SE (HC3) | 95% CI | *p* value |
| --- | --- | --- | --- | --- | --- |
| Total PLEs  - Total score | Birth status | 1.199 | 0.606 | 0.011; 2.388 | 0.049* |
|  | Age | -0.506 | 0.157 | -0.815; -0.198 | 0.001* |
|  | Income decile | -0.244 | 0.081 | -0.402; -0.086 | 0.003* |
|  | Sex | -0.039 | 0.498 | -1.016; 0.938 | 0.938 |
|  | SDQ - Total | 0.131 | 0.044 | 0.044; 0.218 | 0.004* |
| Total PLEs  - Distressing items score | Birth status | 0.435 | 0.411 | -0.370; 1.241 | 0.291 |
|  | Age | -0.173 | 0.118 | -0.405; 0.059 | 0.145 |
|  | Income decile | -0.128 | 0.056 | -0.239; -0.018 | 0.024* |
|  | Sex | 0.338 | 0.356 | -0.360; 1.035 | 0.344 |
|  | SDQ - Total | 0.101 | 0.037 | 0.028; 0.173 | 0.007* |
| Total PLEs  - Distress severity score | Birth status | 1.195 | 1.316 | -1.385; 3.775 | 0.365 |
|  | Age | -0.380 | 0.458 | -1.277; 0.517 | 0.407 |
|  | Income decile | -0.418 | 0.183 | -0.777; -0.060 | 0.023* |
|  | Sex | 1.130 | 1.194 | -1.211; 3.470 | 0.345 |
|  | SDQ - Total | 0.328 | 0.127 | 0.078; 0.577 | 0.011* |
| Unusual thought content  - Total score | Birth status | 0.472 | 0.304 | -0.124; 1.068 | 0.122 |
|  | Age | -0.219 | 0.077 | -0.370; -0.068 | 0.005* |
|  | Income decile | -0.077 | 0.040 | -0.155; 0.000 | 0.052 |
|  | Sex | 0.142 | 0.250 | -0.348; 0.631 | 0.571 |
|  | SDQ - Total | 0.042 | 0.022 | -0.001; 0.085 | 0.056 |
| Unusual thought content  - Distressing items score | Birth status | 0.156 | 0.196 | -0.227; 0.540 | 0.425 |
|  | Age | -0.059 | 0.052 | -0.160; 0.042 | 0.257 |
|  | Income decile | -0.042 | 0.026 | -0.094; 0.009 | 0.111 |
|  | Sex | 0.185 | 0.172 | -0.153; 0.523 | 0.284 |
|  | SDQ - Total | 0.031 | 0.017 | -0.002; 0.064 | 0.067 |
| Unusual thought content  - Distress severity score | Birth status | 0.487 | 0.683 | -0.851; 1.825 | 0.476 |
|  | Age | -0.165 | 0.197 | -0.551; 0.221 | 0.402 |
|  | Income decile | -0.146 | 0.085 | -0.313; 0.022 | 0.090 |
|  | Sex | 0.574 | 0.600 | -0.603; 1.750 | 0.340 |
|  | SDQ - Total | 0.093 | 0.059 | -0.023; 0.209 | 0.119 |
| Perceptual abnormalities  - Total score | Birth status | 0.645 | 0.263 | 0.130; 1.160 | 0.015* |
|  | Age | -0.213 | 0.071 | -0.352; -0.075 | 0.003* |
|  | Income decile | -0.094 | 0.036 | -0.164; -0.023 | 0.010* |
|  | Sex | -0.131 | 0.216 | -0.553; 0.292 | 0.545 |
|  | SDQ - Total | 0.059 | 0.020 | 0.021; 0.098 | 0.003* |
| Perceptual abnormalities  - Distressing items score | Birth status | 0.366 | 0.181 | 0.010; 0.721 | 0.045* |
|  | Age | -0.088 | 0.055 | -0.196; 0.020 | 0.112 |
|  | Income decile | -0.053 | 0.027 | -0.105; 0.000 | 0.052 |
|  | Sex | 0.051 | 0.155 | -0.252; 0.355 | 0.741 |
|  | SDQ - Total | 0.042 | 0.017 | 0.009; 0.075 | 0.013* |
| Perceptual abnormalities  - Distress severity score | Birth status | 1.010 | 0.515 | 0.001; 2.019 | 0.051 |
|  | Age | -0.186 | 0.193 | -0.564; 0.192 | 0.335 |
|  | Income decile | -0.165 | 0.085 | -0.332; 0.002 | 0.054 |
|  | Sex | 0.201 | 0.473 | -0.725; 1.127 | 0.671 |
|  | SDQ - Total | 0.134 | 0.054 | 0.028; 0.240 | 0.014* |
| Disorganized speech  - Total score | Birth status | 0.047 | 0.108 | -0.165; 0.258 | 0.666 |
|  | Age | -0.038 | 0.029 | -0.095; 0.020 | 0.198 |
|  | Income decile | -0.054 | 0.016 | -0.085; -0.023 | <0.001* |
|  | Sex | -0.043 | 0.086 | -0.212; 0.126 | 0.619 |
|  | SDQ - Total | 0.023 | 0.009 | 0.006; 0.040 | 0.008* |
| Disorganized speech  - Distressing items score | Birth status | -0.058 | 0.084 | -0.223; 0.106 | 0.488 |
|  | Age | -0.006 | 0.024 | -0.054; 0.041 | 0.792 |
|  | Income decile | -0.016 | 0.012 | -0.040; 0.007 | 0.175 |
|  | Sex | 0.062 | 0.067 | -0.070; 0.193 | 0.357 |
|  | SDQ - Total | 0.021 | 0.008 | 0.007; 0.036 | 0.005* |
| Disorganized speech  - Distress severity score | Birth status | -0.051 | 0.247 | -0.534; 0.433 | 0.838 |
|  | Age | -0.009 | 0.085 | -0.176; 0.158 | 0.917 |
|  | Income decile | -0.046 | 0.038 | -0.120; 0.027 | 0.220 |
|  | Sex | 0.212 | 0.211 | -0.202; 0.626 | 0.316 |
|  | SDQ - Total | 0.073 | 0.024 | 0.025; 0.120 | 0.003 |

CI Confidence Interval; HC3 Heteroscedasticity-consistent covariance matrix estimator; SDQ Strengths & Difficulties Questionnaire; SE Standard Error. Birth status was coded as 0 = full-term birth and 1 = very preterm birth. Positive coefficients for Birth status therefore indicate higher symptom scores in the very preterm group relative to the full-term group.

### **eResults 4**. Adjusted regression models of psychotic-like experiences in childhood in the BIPP cohort, additionally adjusted for SDQ – Internalizing problems score

| Outcome | Predictor | ß | Robust SE (HC3) | 95% CI | *p* value |
| --- | --- | --- | --- | --- | --- |
| Total PLEs  - Total score | Birth status | 1.244 | 0.601 | 0.065; 2.422 | 0.040* |
|  | Age | -0.540 | 0.156 | -0.845; -0.234 | <0.001* |
|  | Income decile | -0.260 | 0.081 | -0.419; 0.101 | 0.002* |
|  | Sex | -0.224 | 0.508 | -1.221; 0.772 | 0.660 |
|  | SDQ - Internalizing | 0.212 | 0.078 | 0.060; 0.364 | 0.007* |
| Total PLEs  - Distressing items score | Birth status | 0.423 | 0.411 | -0.382; 1.228 | 0.304 |
|  | Age | -0.195 | 0.116 | -0.424; 0.033 | 0.095 |
|  | Income decile | -0.139 | 0.056 | -0.249; -0.029 | 0.014* |
|  | Sex | 0.167 | 0.369 | -0.556; 0.891 | 0.651 |
|  | SDQ - Internalizing | 0.189 | 0.063 | 0.067; 0.312 | 0.003* |
| Total PLEs  - Distress severity score | Birth status | 1.197 | 1.287 | -1.325; 3.720 | 0.353 |
|  | Age | -0.456 | 0.448 | -1.334; 0.421 | 0.309 |
|  | Income decile | -0.454 | 0.182 | -0.811; -0.097 | 0.013* |
|  | Sex | 0.601 | 1.210 | -1.770; 2.972 | 0.620 |
|  | SDQ - Internalizing | 0.592 | 0.197 | 0.207; 0.978 | 0.003* |
| Unusual thought content  - Total score | Birth status | 0.482 | 0.303 | -0.111; 1.077 | 0.112 |
|  | Age | -0.229 | 0.076 | -0.379; -0.080 | 0.003* |
|  | Income decile | -0.082 | 0.040 | -0.160; -0.004 | 0.039* |
|  | Sex | 0.081 | 0.257 | -0.423; 0.584 | 0.753 |
|  | SDQ - Internalizing | 0.070 | 0.039 | -0.007; 0.146 | 0.075 |
| Unusual thought content  - Distressing items score | Birth status | 0.146 | 0.198 | -0.241; 0.534 | 0.459 |
|  | Age | -0.065 | 0.051 | -0.165; 0.035 | 0.203 |
|  | Income decile | -0.045 | 0.026 | -0.097; 0.007 | 0.089 |
|  | Sex | 0.129 | 0.181 | -0.226; 0.484 | 0.478 |
|  | SDQ - Internalizing | 0.062 | 0.029 | 0.005; 0.119 | 0.035* |
| Unusual thought content  - Distress severity score | Birth status | 0.458 | 0.676 | -0.866; 1.782 | 0.499 |
|  | Age | -0.185 | 0.193 | -0.563; 0.193 | 0.339 |
|  | Income decile | -0.154 | 0.085 | -0.321; 0.012 | 0.071 |
|  | Sex | 0.406 | 0.626 | -0.821; 1.633 | 0.518 |
|  | SDQ - Internalizing | 0.185 | 0.092 | 0.005; 0.365 | 0.045* |
| Perceptual abnormalities  - Total score | Birth status | 0.665 | 0.258 | 0.159; 1.171 | 0.011* |
|  | Age | -0.228 | 0.069 | -0.364; -0.093 | 0.001* |
|  | Income decile | -0.101 | 0.036 | -0.172; -0.030 | 0.006* |
|  | Sex | -0.215 | 0.215 | -0.635; 0.206 | 0.318 |
|  | SDQ - Internalizing | 0.096 | 0.035 | 0.028; 0.164 | 0.006* |
| Perceptual abnormalities  - Distressing items score | Birth status | 0.356 | 0.178 | 0.007; 0.704 | 0.047* |
|  | Age | -0.097 | 0.053 | -0.202; 0.008 | 0.071 |
|  | Income decile | -0.057 | 0.026 | -0.108; -0.005 | 0.032* |
|  | Sex | -0.023 | 0.156 | -0.328; 0.282 | 0.882 |
|  | SDQ - Internalizing | 0.082 | 0.029 | 0.025; 0.140 | 0.006* |
| Perceptual abnormalities  - Distress severity score | Birth status | 1.045 | 0.500 | 0.065; 2.025 | 0.038* |
|  | Age | -0.220 | 0.187 | -0.586; 0.147 | 0.241 |
|  | Income decile | -0.181 | 0.084 | -0.346; -0.016 | 0.033* |
|  | Sex | 0.005 | 0.468 | -0.913; 0.922 | 0.992 |
|  | SDQ - Internalizing | 0.223 | 0.087 | 0.052; 0.395 | 0.011* |
| Disorganized speech  - Total score | Birth status | 0.054 | 0.107 | -0.155; 0.264 | 0.612 |
|  | Age | -0.043 | 0.029 | -0.100; 0.013 | 0.134 |
|  | Income decile | -0.056 | 0.016 | -0.087; -0.026 | <0.001* |
|  | Sex | -0.076 | 0.089 | -0.249; 0.098 | 0.395 |
|  | SDQ - Internalizing | 0.037 | 0.015 | 0.008; 0.066 | 0.013* |
| Disorganized speech  - Distressing items score | Birth status | -0.055 | 0.083 | -0.217; 0.107 | 0.508 |
|  | Age | -0.012 | 0.024 | -0.058; 0.035 | 0.627 |
|  | Income decile | -0.019 | 0.012 | -0.042; 0.005 | 0.120 |
|  | Sex | 0.029 | 0.067 | -0.102; 0.160 | 0.661 |
|  | SDQ - Internalizing | 0.037 | 0.012 | 0.013; 0.061 | 0.003* |
| Disorganized speech  - Distress severity score | Birth status | -0.054 | 0.235 | -0.514; 0.406 | 0.818 |
|  | Age | -0.026 | 0.084 | -0.189; 0.138 | 0.761 |
|  | Income decile | -0.054 | 0.037 | -0.127; 0.019 | 0.151 |
|  | Sex | 0.092 | 0.207 | -0.314; 0.499 | 0.657 |
|  | SDQ - Internalizing | 0.134 | 0.042 | 0.052; 0.216 | 0.002* |

CI Confidence Interval; HC3 Heteroscedasticity-consistent covariance matrix estimator; SDQ Strengths & Difficulties Questionnaire; SE Standard Error. Birth status was coded as 0 = full-term birth and 1 = very preterm birth. Positive coefficients for Birth status therefore indicate higher symptom scores in the very preterm group relative to the full-term group

### **eResults 5**. Adjusted regression models of psychotic-like experiences in childhood in the BIPP cohort, additionally adjusted for SDQ – Externalizing problems score

| Outcome | Predictor | ß | Robust SE (HC3) | 95% CI | *p* value |
| --- | --- | --- | --- | --- | --- |
| Total PLEs  - Total score | Birth status | 1.367 | 0.592 | 0.207; 2.526 | 0.022* |
|  | Age | -0.508 | 0.160 | -0.820; -0.195 | 0.002* |
|  | Income decile | -0.246 | 0.081 | -0.405; -0.088 | 0.003* |
|  | Sex | 0.132 | 0.502 | -0.852; 1.116 | 0.793 |
|  | SDQ - Externalizing | 0.174 | 0.075 | 0.028; 0.321 | 0.021* |
| Total PLEs  - Distressing items score | Birth status | 0.596 | 0.395 | -0.179; 1.371 | 0.133 |
|  | Age | -0.181 | 0.121 | -0.419; 0.056 | 0.136 |
|  | Income decile | -0.134 | 0.057 | -0.245; -0.023 | 0.019* |
|  | Sex | 0.452 | 0.354 | -0.242; 1.146 | 0.203 |
|  | SDQ - Externalizing | 0.112 | 0.059 | -0.003; 0.226 | 0.058 |
| Total PLEs  - Distress severity score | Birth status | 1.690 | 1.285 | -0.829; 4.208 | 0.190 |
|  | Age | -0.401 | 0.467 | -1.316; 0.513 | 0.391 |
|  | Income decile | -0.433 | 0.184 | -0.795; -0.071 | 0.020* |
|  | Sex | 1.517 | 1.186 | -0.807; 3.842 | 0.202 |
|  | SDQ - Externalizing | 0.383 | 0.213 | -0.034; 0.800 | 0.072 |
| Unusual thought content  - Total score | Birth status | 0.527 | 0.297 | -0.055; 1.110 | 0.077 |
|  | Age | -0.220 | 0.078 | -0.372; -0.067 | 0.005* |
|  | Income decile | -0.078 | 0.040 | -0.156; -0.000 | 0.050 |
|  | Sex | 0.195 | 0.251 | -0.296; 0.687 | 0.436 |
|  | SDQ - Externalizing | 0.054 | 0.036 | -0.017; 0.125 | 0.134 |
| Unusual thought content  - Distressing items score | Birth status | 0.210 | 0.188 | -0.157; 0.578 | 0.263 |
|  | Age | -0.062 | 0.053 | -0.165; 0.041 | 0.237 |
|  | Income decile | -0.044 | 0.026 | -0.096; 0.007 | 0.095 |
|  | Sex | 0.218 | 0.167 | -0.109; 0.545 | 0.192 |
|  | SDQ - Externalizing | 0.031 | 0.027 | -0.021; 0.084 | 0.242 |
| Unusual thought content  - Distress severity score | Birth status | 0.649 | 0.662 | -0.649; 1.947 | 0.328 |
|  | Age | -0.176 | 0.200 | -0.569; 0.217 | 0.381 |
|  | Income decile | -0.152 | 0.086 | -0.320; 0.016 | 0.078 |
|  | Sex | 0.672 | 0.569 | -0.442; 1.787 | 0.238 |
|  | SDQ - Externalizing | 0.094 | 0.102 | -0.106; 0.294 | 0.359 |
| Perceptual abnormalities  - Total score | Birth status | 0.721 | 0.258 | 0.216; 1.227 | 0.006* |
|  | Age | -0.214 | 0.072 | -0.355; -0.073 | 0.003* |
|  | Income decile | -0.095 | 0.036 | -0.165; -0.024 | 0.009* |
|  | Sex | -0.053 | 0.220 | -0.484; 0.377 | 0.808 |
|  | SDQ - Externalizing | 0.079 | 0.033 | 0.014; 0.144 | 0.018* |
| Perceptual abnormalities  - Distressing items score | Birth status | 0.437 | 0.175 | 0.093; 0.780 | 0.013* |
|  | Age | -0.092 | 0.057 | -0.204; 0.019 | 0.106 |
|  | Income decile | -0.055 | 0.027 | -0.109; -0.002 | 0.044* |
|  | Sex | 0.097 | 0.158 | -0.212; 0.407 | 0.539 |
|  | SDQ - Externalizing | 0.044 | 0.026 | -0.007; 0.096 | 0.094 |
| Perceptual abnormalities  - Distress severity score | Birth status | 1.189 | 0.503 | 0.203; 2.175 | 0.019* |
|  | Age | -0.189 | 0.197 | -0.575; 0.196 | 0.337 |
|  | Income decile | -0.168 | 0.085 | -0.336; -0.000 | 0.051 |
|  | Sex | 0.373 | 0.480 | -0.568; 1.313 | 0.438 |
|  | SDQ - Externalizing | 0.174 | 0.087 | 0.002; 0.345 | 0.048* |
| Disorganized speech  - Total score | Birth status | 0.076 | 0.107 | -0.134; 0.286 | 0.479 |
|  | Age | -0.038 | 0.029 | -0.095; 0.019 | 0.194 |
|  | Income decile | -0.054 | 0.016 | -0.085; -0.023 | <0.001* |
|  | Sex | -0.013 | 0.088 | -0.185; 0.160 | 0.884 |
|  | SDQ - Externalizing | 0.031 | 0.014 | 0.003; 0.058 | 0.029* |
| Disorganized speech  - Distressing items score | Birth status | -0.028 | 0.085 | -0.194; 0.137 | 0.739 |
|  | Age | -0.007 | 0.025 | -0.056; 0.041 | 0.771 |
|  | Income decile | -0.017 | 0.012 | -0.040; 0.006 | 0.155 |
|  | Sex | 0.088 | 0.072 | -0.052; 0.229 | 0.220 |
|  | SDQ - Externalizing | 0.027 | 0.012 | 0.002; 0.051 | 0.033* |
| Disorganized speech  - Distress severity score | Birth status | 0.062 | 0.255 | -0.437; 0.561 | 0.807 |
|  | Age | -0.014 | 0.088 | -0.186; 0.158 | 0.872 |
|  | Income decile | -0.050 | 0.037 | -0.123; 0.023 | 0.184 |
|  | Sex | 0.297 | 0.234 | -0.161; 0.74 | 0.205 |
|  | SDQ - Externalizing | 0.083 | 0.040 | 0.004; 0.162 | 0.041* |

CI Confidence Interval; HC3 Heteroscedasticity-consistent covariance matrix estimator; SDQ Strengths & Difficulties Questionnaire; SE Standard Error. Birth status was coded as 0 = full-term birth and 1 = very preterm birth. Positive coefficients for Birth status therefore indicate higher symptom scores in the very preterm group relative to the full-term group

### **eResults 6**. Adjusted regression models of psychotic-like experiences in childhood in the BIPP cohort, additionally adjusted for Cognitive Ability and SDQ – Total difficulties score

| Outcome | Predictor | ß | Robust SE (HC3) | 95% CI | *p* value |
| --- | --- | --- | --- | --- | --- |
| Total PLEs  - Total score | Birth status | 0.909 | 0.611 | -0.287; 2.106 | 0.138 |
|  | Age | -0.574 | 0.169 | -0.905; -0.244 | <0.001* |
|  | Income decile | -0.234 | 0.082 | -0.395; -0.074 | 0.005* |
|  | Sex | -0.196 | 0.518 | -1.212; 0.820 | 0.706 |
|  | Cognitive Ability | -0.034 | 0.018 | -0.069; 0.002 | 0.065 |
|  | SDQ - Total | 0.110 | 0.048 | 0.015; 0.204 | 0.024* |
| Total PLEs  - Distressing items score | Birth status | 0.332 | 0.433 | -0.517; 1.181 | 0.445 |
|  | Age | -0.215 | 0.129 | -0.468; 0.039 | 0.098 |
|  | Income decile | -0.125 | 0.057 | -0.237; -0.014 | 0.028* |
|  | Sex | 0.247 | 0.382 | -0.501; 0.995 | 0.518 |
|  | Cognitive Ability | -0.017 | 0.014 | -0.044; 0.010 | 0.221 |
|  | SDQ - Total | 0.087 | 0.040 | 0.008; 0.166 | 0.031* |
| Total PLEs  - Distress severity score | Birth status | 0.916 | 1.367 | -1.762; 3.595 | 0.503 |
|  | Age | -0.520 | 0.474 | -1.449; 0.410 | 0.274 |
|  | Income decile | -0.398 | 0.187 | -0.764; -0.033 | 0.034* |
|  | Sex | 0.872 | 1.262 | -1.602; 3.346 | 0.490 |
|  | Cognitive Ability | -0.059 | 0.046 | -0.149; 0.031 | 0.203 |
|  | SDQ - Total | 0.277 | 0.143 | -0.004; 0.558 | 0.054 |
| Unusual thought content  - Total score | Birth status | 0.334 | 0.309 | -0.271; 0.939 | 0.281 |
|  | Age | -0.239 | 0.082 | -0.399; -0.078 | 0.004* |
|  | Income decile | -0.074 | 0.041 | -0.153; 0.006 | 0.071 |
|  | Sex | 0.083 | 0.261 | -0.428; 0.595 | 0.750 |
|  | Cognitive Ability | -0.012 | 0.009 | -0.030; 0.007 | 0.218 |
|  | SDQ - Total | 0.037 | 0.024 | -0.010; 0.084 | 0.122 |
| Unusual thought content  - Distressing items score | Birth status | 0.092 | 0.208 | -0.315; 0.500 | 0.657 |
|  | Age | -0.076 | 0.056 | -0.185; 0.034 | 0.178 |
|  | Income decile | -0.042 | 0.027 | -0.094; 0.010 | 0.118 |
|  | Sex | 0.141 | 0.181 | -0.214; 0.496 | 0.437 |
|  | Cognitive Ability | -0.007 | 0.007 | -0.020; 0.006 | 0.281 |
|  | SDQ - Total | 0.027 | 0.018 | -0.009; 0.063 | 0.141 |
| Unusual thought content  - Distress severity score | Birth status | 0.315 | 0.722 | -1.100; 1.730 | 0.663 |
|  | Age | -0.208 | 0.204 | -0.608; 0.193 | 0.310 |
|  | Income decile | -0.149 | 0.087 | -0.319; 0.020 | 0.086 |
|  | Sex | 0.445 | 0.624 | -0.777; 1.667 | 0.476 |
|  | Cognitive Ability | -0.018 | 0.024 | -0.065; 0.030 | 0.467 |
|  | SDQ - Total | 0.083 | 0.067 | -0.049; 0.216 | 0.216 |
| Perceptual abnormalities  - Total score | Birth status | 0.552 | 0.262 | 0.038; 1.067 | 0.036* |
|  | Age | -0.228 | 0.076 | -0.376; -0.080 | 0.003* |
|  | Income decile | -0.096 | 0.037 | -0.168; -0.024 | 0.009* |
|  | Sex | -0.164 | 0.222 | -0.601; 0.272 | 0.460 |
|  | Cognitive Ability | -0.009 | 0.007 | -0.024; 0.005 | 0.214 |
|  | SDQ - Total | 0.053 | 0.021 | 0.012; 0.095 | 0.012* |
| Perceptual abnormalities  - Distressing items score | Birth status | 0.347 | 0.187 | -0.020; 0.713 | 0.065 |
|  | Age | -0.102 | 0.062 | -0.223; 0.019 | 0.100 |
|  | Income decile | -0.054 | 0.027 | -0.106; -0.002 | 0.045* |
|  | Sex | 0.023 | 0.167 | -0.303; 0.350 | 0.889 |
|  | Cognitive Ability | -0.006 | 0.006 | -0.018; 0.007 | 0.376 |
|  | SDQ - Total | 0.037 | 0.019 | -0.000; 0.073 | 0.052 |
| Perceptual abnormalities  - Distress severity score | Birth status | 0.954 | 0.519 | -0.064; 1.971 | 0.067 |
|  | Age | -0.242 | 0.205 | -0.644; 0.160 | 0.239 |
|  | Income decile | -0.160 | 0.085 | -0.326; 0.007 | 0.061 |
|  | Sex | 0.115 | 0.500 | -0.865; 1.095 | 0.818 |
|  | Cognitive Ability | -0.023 | 0.018 | -0.058; 0.012 | 0.195 |
|  | SDQ - Total | 0.111 | 0.060 | -0.007; 0.228 | 0.066 |
| Disorganized speech  - Total score | Birth status | -0.006 | 0.109 | -0.219; 0.207 | 0.956 |
|  | Age | -0.059 | 0.031 | -0.120; 0.002 | 0.059 |
|  | Income decile | -0.049 | 0.016 | -0.080; -0.018 | 0.002* |
|  | Sex | -0.094 | 0.089 | -0.269; 0.081 | 0.294 |
|  | Cognitive Ability | -0.008 | 0.003 | -0.014; -0.001 | 0.017* |
|  | SDQ - Total | 0.017 | 0.009 | -0.001; 0.035 | 0.061 |
| Disorganized speech  - Distressing items score | Birth status | -0.080 | 0.087 | -0.250; 0.090 | 0.356 |
|  | Age | -0.011 | 0.027 | -0.063; 0.041 | 0.680 |
|  | Income decile | -0.015 | 0.012 | -0.039; 0.009 | 0.230 |
|  | Sex | 0.047 | 0.073 | -0.096; 0.191 | 0.516 |
|  | Cognitive Ability | -0.002 | 0.003 | -0.007; 0.004 | 0.588 |
|  | SDQ - Total | 0.021 | 0.008 | 0.005; 0.036 | 0.012* |
| Disorganized speech  - Distress severity score | Birth status | -0.131 | 0.247 | -0.615; 0.354 | 0.598 |
|  | Age | -0.030 | 0.095 | -0.216; 0.156 | 0.753 |
|  | Income decile | -0.038 | 0.041 | -0.118; 0.042 | 0.352 |
|  | Sex | 0.167 | 0.237 | -0.297; 0.632 | 0.481 |
|  | Cognitive Ability | -0.009 | 0.011 | -0.029; 0.012 | 0.421 |
|  | SDQ - Total | 0.067 | 0.027 | 0.014; 0.120 | 0.013* |

CI Confidence Interval; HC3 Heteroscedasticity-consistent covariance matrix estimator; SDQ Strengths & Difficulties Questionnaire; SE Standard Error. Birth status was coded as 0 = full-term birth and 1 = very preterm birth. Positive coefficients for Birth status therefore indicate higher symptom scores in the very preterm group relative to the full-term group

### **eResults 7**. Adjusted regression models of psychotic-like experiences in childhood in the BIPP cohort, additionally adjusted for Cognitive Ability and SDQ – Internalizing difficulties score

| Outcome | Predictor | ß | Robust SE (HC3) | 95% CI | *p* value |
| --- | --- | --- | --- | --- | --- |
| Total PLEs  - Total score | Birth status | 0.939 | 0.606 | -0.249; 2.127 | 0.123 |
|  | Age | -0.611 | 0.164 | -0.932; -0.291 | <0.001* |
|  | Income decile | -0.245 | 0.082 | -0.406; -0.084 | 0.003* |
|  | Sex | -0.353 | 0.523 | -1.377; 0.672 | 0.500 |
|  | Cognitive Ability | -0.036 | 0.018 | -0.071; -0.001 | 0.042* |
|  | SDQ - Inter | 0.169 | 0.082 | 0.008; 0.331 | 0.041* |
| Total PLEs  - Distressing items score | Birth status | 0.315 | 0.433 | -0.533; 1.163 | 0.468 |
|  | Age | -0.235 | 0.124 | -0.479; 0.009 | 0.060 |
|  | Income decile | -0.134 | 0.056 | -0.244; -0.024 | 0.018* |
|  | Sex | 0.096 | 0.390 | -0.668; 0.860 | 0.806 |
|  | Cognitive Ability | -0.017 | 0.013 | -0.043; 0.009 | 0.205 |
|  | SDQ - Inter | 0.167 | 0.065 | 0.038; 0.295 | 0.011* |
| Total PLEs  - Distress severity score | Birth status | 0.892 | 1.340 | -1.735; 3.520 | 0.506 |
|  | Age | -0.592 | 0.455 | -1.483; 0.299 | 0.194 |
|  | Income decile | -0.425 | 0.186 | -0.789; -0.062 | 0.023* |
|  | Sex | 0.412 | 1.265 | -2.067; 2.890 | 0.745 |
|  | Cognitive Ability | -0.060 | 0.044 | -0.147; 0.027 | 0.179 |
|  | SDQ - Inter | 0.506 | 0.217 | 0.081; 0.931 | 0.020* |
| Unusual thought content  - Total score | Birth status | 0.342 | 0.308 | -0.262; 0.947 | 0.268 |
|  | Age | -0.251 | 0.080 | -0.407; -0.094 | 0.002* |
|  | Income decile | -0.077 | 0.041 | -0.157; 0.003 | 0.059 |
|  | Sex | 0.029 | 0.265 | -0.490; 0.549 | 0.912 |
|  | Cognitive Ability | -0.013 | 0.009 | -0.031; 0.006 | 0.175 |
|  | SDQ - Inter | 0.058 | 0.041 | -0.023; 0.140 | 0.160 |
| Unusual thought content  - Distressing items score | Birth status | 0.082 | 0.210 | -0.329; 0.493 | 0.696 |
|  | Age | -0.081 | 0.054 | -0.188; 0.026 | 0.139 |
|  | Income decile | -0.044 | 0.027 | -0.096; 0.008 | 0.097 |
|  | Sex | 0.091 | 0.189 | -0.280; 0.462 | 0.632 |
|  | Cognitive Ability | -0.007 | 0.006 | -0.020; 0.006 | 0.286 |
|  | SDQ - Inter | 0.056 | 0.030 | -0.004; 0.115 | 0.068 |
| Unusual thought content  - Distress severity score | Birth status | 0.285 | 0.717 | -1.120; 1.690 | 0.692 |
|  | Age | -0.224 | 0.196 | -0.609; 0.160 | 0.254 |
|  | Income decile | -0.158 | 0.086 | -0.326; 0.011 | 0.069 |
|  | Sex | 0.291 | 0.651 | -0.985; 1.566 | 0.655 |
|  | Cognitive Ability | -0.017 | 0.023 | -0.062; 0.029 | 0.474 |
|  | SDQ - Inter | 0.171 | 0.100 | -0.026; 0.368 | 0.090 |
| Perceptual abnormalities  - Total score | Birth status | 0.565 | 0.259 | 0.058; 1.072 | 0.030* |
|  | Age | -0.246 | 0.073 | -0.388; -0.104 | <0.001* |
|  | Income decile | -0.101 | 0.037 | -0.173; -0.029 | 0.006* |
|  | Sex | -0.241 | 0.220 | -0.672; 0.189 | 0.273 |
|  | Cognitive Ability | -0.011 | 0.007 | -0.025; 0.004 | 0.155 |
|  | SDQ - Inter | 0.083 | 0.037 | 0.011; 0.155 | 0.025* |
| Perceptual abnormalities  - Distressing items score | Birth status | 0.335 | 0.184 | -0.025; 0.695 | 0.070 |
|  | Age | -0.110 | 0.058 | -0.224; 0.004 | 0.061 |
|  | Income decile | -0.057 | 0.026 | -0.108; -0.006 | 0.029* |
|  | Sex | -0.043 | 0.164 | -0.366; 0.279 | 0.793 |
|  | Cognitive Ability | -0.005 | 0.006 | -0.017; 0.007 | 0.394 |
|  | SDQ - Inter | 0.074 | 0.032 | 0.012; 0.136 | 0.021* |
| Perceptual abnormalities  - Distress severity score | Birth status | 0.970 | 0.504 | -0.018; 1.958 | 0.055 |
|  | Age | -0.277 | 0.194 | -0.657; 0.103 | 0.155 |
|  | Income decile | -0.171 | 0.084 | -0.335; -0.006 | 0.043* |
|  | Sex | -0.052 | 0.487 | -1.007; 0.903 | 0.915 |
|  | Cognitive Ability | -0.025 | 0.018 | -0.061; 0.010 | 0.166 |
|  | SDQ - Inter | 0.182 | 0.098 | -0.010; 0.373 | 0.065 |
| Disorganized speech  - Total score | Birth status | -0.001 | 0.108 | -0.213; 0.211 | 0.995 |
|  | Age | -0.065 | 0.030 | -0.125; -0.005 | 0.033* |
|  | Income decile | -0.050 | 0.016 | -0.081; -0.020 | 0.001* |
|  | Sex | -0.118 | 0.090 | -0.295; 0.058 | 0.191 |
|  | Cognitive Ability | -0.008 | 0.003 | -0.014; -0.002 | 0.007* |
|  | SDQ - Inter | 0.026 | 0.015 | -0.004; 0.056 | 0.086 |
| Disorganized speech  - Distressing items score | Birth status | -0.078 | 0.086 | -0.247; 0.092 | 0.370 |
|  | Age | -0.017 | 0.025 | -0.067; 0.032 | 0.496 |
|  | Income decile | -0.017 | 0.012 | -0.041; 0.007 | 0.173 |
|  | Sex | 0.016 | 0.071 | -0.122; 0.155 | 0.818 |
|  | Cognitive Ability | -0.002 | 0.003 | -0.007; 0.003 | 0.460 |
|  | SDQ - Inter | 0.034 | 0.013 | 0.009; 0.059 | 0.007* |
| Disorganized speech  - Distress severity score | Birth status | -0.135 | 0.241 | -0.608; 0.338 | 0.576 |
|  | Age | -0.048 | 0.090 | -0.223; 0.128 | 0.595 |
|  | Income decile | -0.045 | 0.041 | -0.125; 0.035 | 0.275 |
|  | Sex | 0.056 | 0.223 | -0.381; 0.493 | 0.802 |
|  | Cognitive Ability | -0.009 | 0.009 | -0.027; 0.009 | 0.342 |
|  | SDQ - Inter | 0.122 | 0.043 | 0.037; 0.207 | 0.005* |

CI Confidence Interval; HC3 Heteroscedasticity-consistent covariance matrix estimator; SDQ Strengths & Difficulties Questionnaire; SE Standard Error. Birth status was coded as 0 = full-term birth and 1 = very preterm birth. Positive coefficients for Birth status therefore indicate higher symptom scores in the very preterm group relative to the full-term group

### **eResults 8**. Adjusted regression models of psychotic-like experiences in childhood in the BIPP cohort, additionally adjusted for Cognitive Ability and SDQ – Externalizing difficulties score

| Outcome | Predictor | ß | Robust SE (HC3) | 95% CI | *p* value |
| --- | --- | --- | --- | --- | --- |
| Total PLEs  - Total score | Birth status | 1.014 | 0.602 | -0.167; 2.195 | 0.094 |
|  | Age | -0.587 | 0.170 | -0.921; -0.253 | <0.001* |
|  | Income decile | -0.231 | 0.082 | -0.391; -0.070 | 0.005* |
|  | Sex | -0.071 | 0.526 | -1.102; 0.960 | 0.893 |
|  | Cognitive Ability | -0.040 | 0.018 | -0.075; -0.005 | 0.026* |
|  | SDQ - Exter | 0.145 | 0.080 | -0.011; 0.301 | 0.071 |
| Total PLEs  - Distressing items score | Birth status | 0.440 | 0.426 | -0.394; 1.275 | 0.302 |
|  | Age | -0.238 | 0.132 | -0.498; 0.021 | 0.073 |
|  | Income decile | -0.126 | 0.057 | -0.237; -0.014 | 0.028* |
|  | Sex | 0.319 | 0.384 | -0.434; 1.073 | 0.407 |
|  | Cognitive Ability | -0.024 | 0.014 | -0.052; 0.004 | 0.092 |
|  | SDQ - Exter | 0.087 | 0.064 | -0.038; 0.213 | 0.173 |
| Total PLEs  - Distress severity score | Birth status | 1.242 | 1.357 | -1.417; 3.902 | 0.361 |
|  | Age | -0.584 | 0.480 | -1.524; 0.356 | 0.225 |
|  | Income decile | -0.397 | 0.187 | -0.764; -0.029 | 0.035* |
|  | Sex | 1.123 | 1.262 | -1.350; 3.597 | 0.374 |
|  | Cognitive Ability | -0.079 | 0.044 | -0.166; 0.008 | 0.077 |
|  | SDQ - Exter | 0.299 | 0.232 | -0.156; 0.754 | 0.199 |
| Unusual thought content  - Total score | Birth status | 0.370 | 0.306 | -0.229; 0.969 | 0.227 |
|  | Age | -0.243 | 0.083 | -0.407; -0.080 | 0.004* |
|  | Income decile | -0.073 | 0.041 | -0.152; 0.007 | 0.075 |
|  | Sex | 0.124 | 0.265 | -0.395; 0.643 | 0.639 |
|  | Cognitive Ability | -0.014 | 0.009 | -0.032; 0.004 | 0.137 |
|  | SDQ - Exter | 0.048 | 0.039 | -0.029; 0.124 | 0.225 |
| Unusual thought content  - Distressing items score | Birth status | 0.129 | 0.204 | -0.270; 0.529 | 0.527 |
|  | Age | -0.084 | 0.057 | -0.196; 0.027 | 0.141 |
|  | Income decile | -0.042 | 0.027 | -0.094; 0.010 | 0.115 |
|  | Sex | 0.160 | 0.178 | -0.188; 0.509 | 0.368 |
|  | Cognitive Ability | -0.010 | 0.007 | -0.023; 0.004 | 0.161 |
|  | SDQ - Exter | 0.024 | 0.029 | -0.034; 0.081 | 0.418 |
| Unusual thought content  - Distress severity score | Birth status | 0.428 | 0.714 | -0.972; 1.828 | 0.550 |
|  | Age | -0.234 | 0.208 | -0.642; 0.173 | 0.261 |
|  | Income decile | -0.150 | 0.087 | -0.321; 0.020 | 0.085 |
|  | Sex | 0.506 | 0.597 | -0.665; 1.676 | 0.398 |
|  | Cognitive Ability | -0.025 | 0.024 | -0.071; 0.022 | 0.298 |
|  | SDQ - Exter | 0.075 | 0.113 | -0.147; 0.297 | 0.510 |
| Perceptual abnormalities  - Total score | Birth status | 0.604 | 0.260 | 0.095; 1.113 | 0.021* |
|  | Age | -0.235 | 0.076 | -0.384; -0.085 | 0.002* |
|  | Income decile | -0.095 | 0.037 | -0.166; -0.023 | 0.010* |
|  | Sex | -0.105 | 0.227 | -0.550; 0.340 | 0.645 |
|  | Cognitive Ability | -0.012 | 0.007 | -0.027; 0.002 | 0.089 |
|  | SDQ - Exter | 0.069 | 0.035 | 0.001; 0.138 | 0.048* |
| Perceptual abnormalities  - Distressing items score | Birth status | 0.395 | 0.184 | 0.035; 0.755 | 0.032* |
|  | Age | -0.113 | 0.063 | -0.237; 0.010 | 0.073 |
|  | Income decile | -0.054 | 0.027 | -0.107; -0.001 | 0.046* |
|  | Sex | 0.051 | 0.172 | -0.286; 0.387 | 0.768 |
|  | Cognitive Ability | -0.009 | 0.006 | -0.021; 0.003 | 0.162 |
|  | SDQ - Exter | 0.034 | 0.029 | -0.022; 0.090 | 0.240 |
| Perceptual abnormalities  - Distress severity score | Birth status | 1.068 | 0.516 | 0.056; 2.079 | 0.040* |
|  | Age | -0.260 | 0.205 | -0.662; 0.143 | 0.207 |
|  | Income decile | -0.157 | 0.085 | -0.325; 0.010 | 0.066 |
|  | Sex | 0.232 | 0.509 | -0.766; 1.230 | 0.649 |
|  | Cognitive Ability | -0.030 | 0.016 | -0.062; 0.002 | 0.068 |
|  | SDQ - Exter | 0.137 | 0.093 | -0.046; 0.319 | 0.143 |
| Disorganized speech  - Total score | Birth status | 0.010 | 0.108 | -0.201; 0.221 | 0.925 |
|  | Age | -0.061 | 0.031 | -0.122; -0.000 | 0.050 |
|  | Income decile | -0.048 | 0.016 | -0.079; -0.017 | 0.003* |
|  | Sex | -0.074 | 0.092 | -0.255; 0.107 | 0.425 |
|  | Cognitive Ability | -0.009 | 0.003 | -0.015; -0.002 | 0.008* |
|  | SDQ - Exter | 0.023 | 0.015 | -0.006; 0.052 | 0.115 |
| Disorganized speech  - Distressing items score | Birth status | -0.059 | 0.087 | -0.230; 0.112 | 0.501 |
|  | Age | -0.014 | 0.028 | -0.068; 0.040 | 0.602 |
|  | Income decile | -0.015 | 0.012 | -0.039; 0.010 | 0.240 |
|  | Sex | 0.069 | 0.079 | -0.086; 0.224 | 0.385 |
|  | Cognitive Ability | -0.003 | 0.003 | -0.008; 0.003 | 0.331 |
|  | SDQ - Exter | 0.025 | 0.013 | -0.001; 0.051 | 0.064 |
| Disorganized speech  - Distress severity score | Birth status | -0.052 | 0.252 | -0.546; 0.442 | 0.837 |
|  | Age | -0.045 | 0.099 | -0.239; 0.149 | 0.649 |
|  | Income decile | -0.038 | 0.041 | -0.117; 0.042 | 0.356 |
|  | Sex | 0.229 | 0.265 | -0.291; 0.749 | 0.389 |
|  | Cognitive Ability | -0.013 | 0.011 | -0.035; 0.008 | 0.224 |
|  | SDQ - Exter | 0.074 | 0.045 | -0.015; 0.162 | 0.106 |

CI Confidence Interval; HC3 Heteroscedasticity-consistent covariance matrix estimator; SDQ Strengths & Difficulties Questionnaire; SE Standard Error. Birth status was coded as 0 = full-term birth and 1 = very preterm birth. Positive coefficients for Birth status therefore indicate higher symptom scores in the very preterm group relative to the full-term group

### **eResults 9**. Adjusted regression models of psychotic-like experiences in childhood in the ABCD cohort, additionally adjusted for Cognitive Ability

| Outcome | Predictor | ß | SE | 95% CI | *p* value |
| --- | --- | --- | --- | --- | --- |
| Total PLEs  - Total score | Birth status | -0.288 | 0.299 | -0.874; 0.298 | 0.336 |
|  | Age | -0.273 | 0.055 | -0.380; -0.166 | <0.001* |
|  | Area Deprivation Index | 0.023 | 0.002 | 0.018; 0.027 | <0.001* |
|  | Sex | -0.331 | 0.069 | -0.466; -0.196 | <0.001* |
|  | Cognitive Ability | -0.069 | 0.012 | -0.093; -0.046 | <0.001* |
| Total PLEs  - Distressing items score | Birth status | 0.033 | 0.193 | -0.345; 0.410 | 0.866 |
|  | Age | -0.208 | 0.035 | -0.277; -0.138 | <0.001* |
|  | Area Deprivation Index | 0.012 | 0.001 | 0.009; 0.015 | <0.001* |
|  | Sex | -0.093 | 0.045 | -0.181; -0.006 | 0.037* |
|  | Cognitive Ability | -0.049 | 0.008 | -0.065; -0.034 | <0.001* |
| Total PLEs  - Distress severity score | Birth status | 0.182 | 0.624 | -1.040; 1.405 | 0.770 |
|  | Age | -0.644 | 0.115 | -0.870; -0.418 | <0.001* |
|  | Area Deprivation Index | 0.042 | 0.005 | 0.033; 0.051 | <0.001* |
|  | Sex | -0.107 | 0.145 | -0.392; 0.178 | 0.461 |
|  | Cognitive Ability | -0.182 | 0.025 | -0.231; -0.132 | <0.001* |
| Unusual thought content  - Total score | Birth status | -0.173 | 0.153 | -0.474; 0.127 | 0.258 |
|  | Age | -0.125 | 0.028 | -0.180; -0.070 | <0.001* |
|  | Area Deprivation Index | 0.010 | 0.001 | 0.008; 0.013 | <0.001* |
|  | Sex | -0.073 | 0.036 | -0.143; -0.003 | 0.040* |
|  | Cognitive Ability | -0.029 | 0.006 | -0.041; -0.017 | <0.001* |
| Unusual thought content  - Distressing items score | Birth status | 0.016 | 0.101 | -0.182; 0.213 | 0.876 |
|  | Age | -0.098 | 0.019 | -0.134; -0.061 | <0.001* |
|  | Area Deprivation Index | 0.005 | 0.001 | 0.004; 0.007 | <0.001* |
|  | Sex | 0.018 | 0.024 | -0.028; 0.065 | 0.436 |
|  | Cognitive Ability | -0.021 | 0.004 | -0.029; -0.013 | <0.001* |
| Unusual thought content  - Distress severity score | Birth status | 0.041 | 0.339 | -0.622; 0.705 | 0.903 |
|  | Age | -0.326 | 0.063 | -0.449; -0.203 | <0.001* |
|  | Area Deprivation Index | 0.019 | 0.003 | 0.014; 0.024 | <0.001* |
|  | Sex | 0.151 | 0.079 | -0.003; 0.306 | 0.055 |
|  | Cognitive Ability | -0.084 | 0.014 | -0.111; -0.057 | <0.001* |
| Perceptual abnormalities  - Total score | Birth status | -0.100 | 0.139 | -0.372; 0.171 | 0.469 |
|  | Age | -0.130 | 0.026 | -0.180; -0.080 | <0.001* |
|  | Area Deprivation Index | 0.009 | 0.001 | 0.007; 0.011 | <0.001* |
|  | Sex | -0.196 | 0.032 | -0.259; -0.133 | <0.001* |
|  | Cognitive Ability | -0.026 | 0.006 | -0.037; -0.015 | <0.001* |
| Perceptual abnormalities  - Distressing items score | Birth status | 0.009 | 0.093 | -0.175; 0.192 | 0.927 |
|  | Age | -0.103 | 0.017 | -0.137; -0.069 | <0.001* |
|  | Area Deprivation Index | 0.006 | 0.001 | 0.004; 0.007 | <0.001* |
|  | Sex | -0.084 | 0.022 | -0.126; -0.041 | <0.001* |
|  | Cognitive Ability | -0.022 | 0.004 | -0.030; -0.015 | <0.001* |
| Perceptual abnormalities  - Distress severity score | Birth status | 0.127 | 0.290 | -0.441; 0.696 | 0.661 |
|  | Age | -0.302 | 0.054 | -0.408; -0.196 | <0.001* |
|  | Area Deprivation Index | 0.019 | 0.002 | 0.015; 0.024 | <0.001* |
|  | Sex | -0.200 | 0.068 | -0.334; -0.067 | 0.003* |
|  | Cognitive Ability | -0.079 | 0.012 | -0.103; -0.056 | <0.001* |
| Disorganized speech  - Total score | Birth status | -0.015 | 0.054 | -0.120; 0.091 | 0.782 |
|  | Age | -0.019 | 0.010 | -0.039; 0.001 | 0.060 |
|  | Area Deprivation Index | 0.004 | 0.000 | 0.003; 0.004 | <0.001* |
|  | Sex | -0.059 | 0.013 | -0.084; -0.034 | <0.001* |
|  | Cognitive Ability | -0.015 | 0.002 | -0.019; -0.010 | <0.001* |
| Disorganized speech  - Distressing items score | Birth status | 0.006 | 0.034 | -0.061; 0.074 | 0.851 |
|  | Age | -0.006 | 0.007 | -0.019; 0.007 | 0.343 |
|  | Area Deprivation Index | 0.001 | 0.000 | 0.000; 0.001 | 0.003* |
|  | Sex | -0.026 | 0.008 | -0.042; -0.010 | 0.002* |
|  | Cognitive Ability | -0.006 | 0.001 | -0.009; -0.003 | <0.001* |
| Disorganized speech  - Distress severity score | Birth status | 0.013 | 0.092 | -0.167; 0.192 | 0.890 |
|  | Age | -0.013 | 0.017 | -0.047; 0.021 | 0.450 |
|  | Area Deprivation Index | 0.002 | 0.001 | 0.001; 0.004 | <0.001* |
|  | Sex | -0.054 | 0.022 | -0.097; -0.011 | 0.013* |
|  | Cognitive Ability | -0.019 | 0.004 | -0.027; -0.012 | <0.001* |

CI Confidence Interval; SE Standard Error. Birth status was coded as 0 = full-term birth and 1 = very preterm birth. Positive coefficients for Birth status therefore indicate higher symptom scores in the very preterm group relative to the full-term group

### **eResults 10**. Adjusted regression models of psychotic-like experiences in childhood in the ABCD cohort, additionally adjusted for CBCL – Total problem score

| Outcome | Predictor | ß | SE | 95% CI | *p* value |
| --- | --- | --- | --- | --- | --- |
| Total PLEs  - Total score | Birth status | -0.231 | 0.294 | -0.808; 0.346 | 0.432 |
|  | Age | -0.272 | 0.054 | -0.377; -0.167 | <0.001* |
|  | Area Deprivation Index | 0.023 | 0.002 | 0.019; 0.028 | <0.001* |
|  | Sex | -0.266 | 0.068 | -0.400; -0.133 | <0.001* |
|  | CBCL- Total | 0.024 | 0.002 | 0.020; 0.028 | <0.001* |
| Total PLEs  - Distressing items score | Birth status | 0.064 | 0.190 | -0.308; 0.435 | 0.736 |
|  | Age | -0.207 | 0.035 | -0.275; -0.139 | <0.001* |
|  | Area Deprivation Index | 0.013 | 0.001 | 0.010; 0.016 | <0.001* |
|  | Sex | -0.053 | 0.044 | -0.139; 0.034 | 0.232 |
|  | CBCL- Total | 0.015 | 0.001 | 0.013; 0.018 | <0.001* |
| Total PLEs  - Distress severity score | Birth status | 0.309 | 0.614 | -0.894; 1.512 | 0.615 |
|  | Age | -0.641 | 0.113 | -0.862; -0.420 | <0.001* |
|  | Area Deprivation Index | 0.045 | 0.005 | 0.036; 0.054 | <0.001* |
|  | Sex | 0.015 | 0.143 | -0.266; 0.295 | 0.917 |
|  | CBCL- Total | 0.050 | 0.004 | 0.042; 0.058 | <0.001* |
| Unusual thought content  - Total score | Birth status | -0.151 | 0.150 | -0.446; 0.143 | 0.315 |
|  | Age | -0.126 | 0.028 | -0.180; -0.071 | <0.001* |
|  | Area Deprivation Index | 0.010 | 0.001 | 0.008; 0.013 | <0.001* |
|  | Sex | -0.034 | 0.035 | -0.103; 0.035 | 0.335 |
|  | CBCL- Total | 0.012 | 0.001 | 0.010; 0.014 | <0.001* |
| Unusual thought content  - Distressing items score | Birth status | 0.022 | 0.099 | -0.173; 0.217 | 0.824 |
|  | Age | -0.097 | 0.018 | -0.133; -0.061 | <0.001* |
|  | Area Deprivation Index | 0.006 | 0.001 | 0.004; 0.007 | <0.001* |
|  | Sex | 0.040 | 0.023 | -0.005; 0.086 | 0.084 |
|  | CBCL- Total | 0.007 | 0.001 | 0.006; 0.009 | <0.001* |
| Unusual thought content  - Distress severity score | Birth status | 0.074 | 0.333 | -0.579; 0.727 | 0.825 |
|  | Age | -0.324 | 0.061 | -0.444; -0.203 | <0.001* |
|  | Area Deprivation Index | 0.021 | 0.002 | 0.016; 0.026 | <0.001* |
|  | Sex | 0.215 | 0.078 | 0.063; 0.368 | 0.006* |
|  | CBCL- Total | 0.025 | 0.002 | 0.021; 0.030 | <0.001* |
| Perceptual abnormalities  - Total score | Birth status | -0.064 | 0.137 | -0.332; 0.205 | 0.642 |
|  | Age | -0.130 | 0.025 | -0.179; -0.081 | <0.001* |
|  | Area Deprivation Index | 0.009 | 0.001 | 0.007; 0.011 | <0.001* |
|  | Sex | -0.173 | 0.032 | -0.235; -0.110 | <0.001* |
|  | CBCL- Total | 0.010 | 0.001 | 0.008; 0.012 | <0.001* |
| Perceptual abnormalities  - Distressing items score | Birth status | 0.034 | 0.092 | -0.146; 0.215 | 0.709 |
|  | Age | -0.103 | 0.017 | -0.136; -0.070 | <0.001* |
|  | Area Deprivation Index | 0.006 | 0.001 | 0.005; 0.007 | <0.001* |
|  | Sex | -0.068 | 0.021 | -0.110; -0.026 | 0.002* |
|  | CBCL- Total | 0.006 | 0.001 | 0.005; 0.008 | <0.001* |
| Perceptual abnormalities  - Distress severity score | Birth status | 0.217 | 0.286 | -0.343; 0.776 | 0.448 |
|  | Age | -0.301 | 0.053 | -0.405; -0.197 | <0.001* |
|  | Area Deprivation Index | 0.020 | 0.002 | 0.016; 0.025 | <0.001* |
|  | Sex | -0.152 | 0.067 | -0.284; -0.021 | 0.023* |
|  | CBCL- Total | 0.021 | 0.002 | 0.017; 0.024 | <0.001* |
| Disorganized speech  - Total score | Birth status | -0.013 | 0.053 | -0.118; 0.091 | 0.801 |
|  | Age | -0.017 | 0.010 | -0.036; 0.003 | 0.094 |
|  | Area Deprivation Index | 0.004 | 0.000 | 0.003; 0.005 | <0.001* |
|  | Sex | -0.057 | 0.013 | -0.082; -0.032 | <0.001* |
|  | CBCL- Total | 0.002 | 0.000 | 0.001; 0.003 | <0.001* |
| Disorganized speech  - Distressing items score | Birth status | 0.007 | 0.034 | -0.060; 0.074 | 0.830 |
|  | Age | -0.006 | 0.006 | -0.019; 0.007 | 0.355 |
|  | Area Deprivation Index | 0.001 | 0.000 | 0.000; 0.001 | <0.001* |
|  | Sex | -0.022 | 0.008 | -0.038; -0.007 | 0.006* |
|  | CBCL- Total | 0.001 | 0.000 | 0.001; 0.002 | <0.001* |
| Disorganized speech  - Distress severity score | Birth status | 0.021 | 0.091 | -0.157; 0.200 | 0.814 |
|  | Age | -0.013 | 0.017 | -0.047; 0.020 | 0.439 |
|  | Area Deprivation Index | 0.003 | 0.001 | 0.002; 0.004 | <0.001* |
|  | Sex | -0.044 | 0.022 | -0.086; -0.002 | 0.042* |
|  | CBCL- Total | 0.004 | 0.001 | 0.003; 0.005 | <0.001* |

CBCL Child Behaviour Checklist; CI Confidence Interval; SE Standard Error. Birth status was coded as 0 = full-term birth and 1 = very preterm birth. Positive coefficients for Birth status therefore indicate higher symptom scores in the very preterm group relative to the full-term group

### **eResults 11**. Adjusted regression models of psychotic-like experiences in childhood in the ABCD cohort, additionally adjusted for CBCL – Internalizing problems score

| Outcome | Predictor | ß | SE | 95% CI | *p* value |
| --- | --- | --- | --- | --- | --- |
| Total PLEs  - Total score | Birth status | -0.271 | 0.296 | -0.851; 0.309 | 0.360 |
|  | Age | -0.283 | 0.054 | -0.389; -0.178 | <0.001* |
|  | Area Deprivation Index | 0.025 | 0.002 | 0.020; 0.029 | <0.001* |
|  | Sex | -0.351 | 0.068 | -0.484; -0.218 | <0.001* |
|  | CBCL- Inter | 0.045 | 0.006 | 0.033; 0.058 | <0.001* |
| Total PLEs  - Distressing items score | Birth status | 0.038 | 0.190 | -0.335; 0.411 | 0.841 |
|  | Age | -0.214 | 0.035 | -0.282; -0.145 | <0.001* |
|  | Area Deprivation Index | 0.014 | 0.001 | 0.011; 0.016 | <0.001* |
|  | Sex | -0.106 | 0.044 | -0.193; -0.020 | 0.016* |
|  | CBCL- Inter | 0.030 | 0.004 | 0.022; 0.038 | <0.001* |
| Total PLEs  - Distress severity score | Birth status | 0.225 | 0.617 | -0.984; 1.434 | 0.715 |
|  | Age | -0.663 | 0.114 | -0.886; -0.440 | <0.001* |
|  | Area Deprivation Index | 0.048 | 0.005 | 0.039; 0.057 | <0.001* |
|  | Sex | -0.162 | 0.143 | -0.443; 0.118 | 0.257 |
|  | CBCL- Inter | 0.095 | 0.013 | 0.069; 0.121 | <0.001* |
| Unusual thought content  - Total score | Birth status | -0.171 | 0.151 | -0.468; 0.125 | 0.256 |
|  | Age | -0.131 | 0.028 | -0.186; -0.077 | <0.001* |
|  | Area Deprivation Index | 0.011 | 0.001 | 0.009; 0.013 | <0.001* |
|  | Sex | -0.077 | 0.035 | -0.146; -0.008 | 0.028* |
|  | CBCL- Inter | 0.023 | 0.003 | 0.017; 0.029 | <0.001* |
| Unusual thought content  - Distressing items score | Birth status | 0.010 | 0.100 | -0.186; 0.205 | 0.923 |
|  | Age | -0.100 | 0.018 | -0.136; -0.064 | <0.001* |
|  | Area Deprivation Index | 0.006 | 0.001 | 0.005; 0.008 | <0.001* |
|  | Sex | 0.014 | 0.023 | -0.032; 0.059 | 0.549 |
|  | CBCL- Inter | 0.014 | 0.002 | 0.010; 0.019 | <0.001* |
| Unusual thought content  - Distress severity score | Birth status | 0.031 | 0.335 | -0.625; 0.687 | 0.926 |
|  | Age | -0.335 | 0.062 | -0.456; -0.214 | <0.001* |
|  | Area Deprivation Index | 0.022 | 0.002 | 0.017; 0.027 | <0.001* |
|  | Sex | 0.125 | 0.078 | -0.027; 0.278 | 0.106 |
|  | CBCL- Inter | 0.049 | 0.007 | 0.035; 0.063 | <0.001* |
| Perceptual abnormalities  - Total score | Birth status | -0.081 | 0.137 | -0.350; 0.188 | 0.557 |
|  | Age | -0.135 | 0.025 | -0.184; -0.085 | <0.001* |
|  | Area Deprivation Index | 0.009 | 0.001 | 0.007; 0.011 | <0.001* |
|  | Sex | -0.208 | 0.032 | -0.270; -0.145 | <0.001* |
|  | CBCL- Inter | 0.020 | 0.003 | 0.015; 0.026 | <0.001* |
| Perceptual abnormalities  - Distressing items score | Birth status | 0.023 | 0.092 | -0.158; 0.204 | 0.803 |
|  | Age | -0.106 | 0.017 | -0.139; -0.073 | <0.001* |
|  | Area Deprivation Index | 0.006 | 0.001 | 0.005; 0.008 | <0.001* |
|  | Sex | -0.091 | 0.021 | -0.133; -0.049 | <0.001* |
|  | CBCL- Inter | 0.014 | 0.002 | 0.010; 0.017 | <0.001* |
| Perceptual abnormalities  - Distress severity score | Birth status | 0.182 | 0.287 | -0.380; 0.743 | 0.526 |
|  | Age | -0.310 | 0.053 | -0.414; -0.205 | <0.001* |
|  | Area Deprivation Index | 0.022 | 0.002 | 0.018; 0.026 | <0.001* |
|  | Sex | -0.225 | 0.067 | -0.357; -0.094 | <0.001* |
|  | CBCL- Inter | 0.039 | 0.006 | 0.027; 0.051 | <0.001* |
| Disorganized speech  - Total score | Birth status | -0.016 | 0.053 | -0.121; 0.089 | 0.763 |
|  | Age | -0.017 | 0.010 | -0.037; 0.002 | 0.084 |
|  | Area Deprivation Index | 0.004 | 0.000 | 0.003; 0.005 | <0.001* |
|  | Sex | -0.063 | 0.013 | -0.088; -0.039 | <0.001* |
|  | CBCL- Inter | 0.002 | 0.001 | -0.000; 0.004 | 0.067 |
| Disorganized speech  - Distressing items score | Birth status | 0.005 | 0.034 | -0.062; 0.072 | 0.878 |
|  | Age | -0.007 | 0.006 | -0.019; 0.006 | 0.316 |
|  | Area Deprivation Index | 0.001 | 0.000 | 0.000; 0.001 | <0.001* |
|  | Sex | -0.027 | 0.008 | -0.043; -0.011 | <0.001* |
|  | CBCL- Inter | 0.002 | 0.001 | 0.001; 0.004 | 0.003* |
| Disorganized speech  - Distress severity score | Birth status | 0.015 | 0.091 | -0.164; 0.193 | 0.870 |
|  | Age | -0.015 | 0.017 | -0.048; 0.019 | 0.385 |
|  | Area Deprivation Index | 0.003 | 0.001 | 0.002; 0.004 | <0.001* |
|  | Sex | -0.058 | 0.021 | -0.100; -0.016 | 0.007* |
|  | CBCL- Inter | 0.007 | 0.002 | 0.003; 0.011 | <0.001* |

CBCL Child Behaviour Checklist; CI Confidence Interval; SE Standard Error. Birth status was coded as 0 = full-term birth and 1 = very preterm birth. Positive coefficients for Birth status therefore indicate higher symptom scores in the very preterm group relative to the full-term group

### **eResults 12**. Adjusted regression models of psychotic-like experiences in childhood in the ABCD cohort, additionally adjusted for CBCL – Externalizing problems score

| Outcome | Predictor | ß | SE | 95% CI | *p* value |
| --- | --- | --- | --- | --- | --- |
| Total PLEs  - Total score | Birth status | -0.216 | 0.295 | -0.795; 0.363 | 0.464 |
|  | Age | -0.268 | 0.054 | -0.373; -0.162 | <0.001* |
|  | Area Deprivation Index | 0.024 | 0.002 | 0.019; 0.028 | <0.001* |
|  | Sex | -0.272 | 0.068 | -0.406; -0.138 | <0.001* |
|  | CBCL- Exter | 0.059 | 0.006 | 0.048; 0.071 | <0.001* |
| Total PLEs  - Distressing items score | Birth status | 0.073 | 0.190 | -0.299; 0.446 | 0.700 |
|  | Age | -0.204 | 0.035 | -0.272; -0.136 | <0.001* |
|  | Area Deprivation Index | 0.013 | 0.001 | 0.010; 0.016 | <0.001* |
|  | Sex | -0.057 | 0.044 | -0.144; 0.030 | 0.199 |
|  | CBCL- Exter | 0.037 | 0.004 | 0.030; 0.045 | <0.001* |
| Total PLEs  - Distress severity score | Birth status | 0.343 | 0.615 | -0.864; 1.549 | 0.578 |
|  | Age | -0.631 | 0.113 | -0.853; -0.410 | <0.001* |
|  | Area Deprivation Index | 0.046 | 0.005 | 0.037; 0.054 | <0.001* |
|  | Sex | 0.007 | 0.144 | -0.275; 0.288 | 0.962 |
|  | CBCL- Exter | 0.128 | 0.013 | 0.103; 0.153 | <0.001* |
| Unusual thought content  - Total score | Birth status | -0.143 | 0.151 | -0.439; 0.152 | 0.341 |
|  | Age | -0.123 | 0.028 | -0.178; -0.069 | <0.001* |
|  | Area Deprivation Index | 0.011 | 0.001 | 0.008; 0.013 | <0.001* |
|  | Sex | -0.037 | 0.035 | -0.106; 0.032 | 0.296 |
|  | CBCL- Exter | 0.030 | 0.003 | 0.024; 0.036 | <0.001* |
| Unusual thought content  - Distressing items score | Birth status | 0.027 | 0.100 | -0.168; 0.222 | 0.788 |
|  | Age | -0.095 | 0.018 | -0.131; -0.059 | <0.001* |
|  | Area Deprivation Index | 0.006 | 0.001 | 0.004; 0.007 | <0.001* |
|  | Sex | 0.038 | 0.023 | -0.007; 0.084 | 0.100 |
|  | CBCL- Exter | 0.019 | 0.002 | 0.014; 0.023 | <0.001* |
| Unusual thought content  - Distress severity score | Birth status | 0.090 | 0.334 | -0.564; 0.745 | 0.787 |
|  | Age | -0.319 | 0.062 | -0.440; -0.198 | <0.001* |
|  | Area Deprivation Index | 0.021 | 0.002 | 0.016; 0.026 | <0.001* |
|  | Sex | 0.211 | 0.078 | 0.058; 0.364 | 0.007* |
|  | CBCL- Exter | 0.065 | 0.007 | 0.051; 0.078 | <0.001* |
| Perceptual abnormalities  - Total score | Birth status | -0.058 | 0.137 | -0.327; 0.211 | 0.673 |
|  | Age | -0.128 | 0.025 | -0.178; -0.079 | <0.001* |
|  | Area Deprivation Index | 0.009 | 0.001 | 0.007; 0.011 | <0.001* |
|  | Sex | -0.176 | 0.032 | -0.239; -0.113 | <0.001* |
|  | CBCL- Exter | 0.024 | 0.003 | 0.018; 0.029 | <0.001* |
| Perceptual abnormalities  - Distressing items score | Birth status | 0.038 | 0.092 | -0.143; 0.219 | 0.680 |
|  | Age | -0.102 | 0.017 | -0.135; -0.068 | <0.001* |
|  | Area Deprivation Index | 0.006 | 0.001 | 0.005; 0.008 | <0.001* |
|  | Sex | -0.070 | 0.022 | -0.112; -0.028 | 0.001* |
|  | CBCL- Exter | 0.016 | 0.002 | 0.012; 0.020 | <0.001* |
| Perceptual abnormalities  - Distress severity score | Birth status | 0.231 | 0.286 | -0.330; 0.791 | 0.420 |
|  | Age | -0.297 | 0.053 | -0.401; -0.193 | <0.001* |
|  | Area Deprivation Index | 0.021 | 0.002 | 0.017; 0.025 | <0.001* |
|  | Sex | -0.155 | 0.067 | -0.287; -0.023 | 0.021* |
|  | CBCL- Exter | 0.053 | 0.006 | 0.041; 0.065 | <0.001* |
| Disorganized speech  - Total score | Birth status | -0.012 | 0.053 | -0.117; 0.093 | 0.823 |
|  | Age | -0.016 | 0.010 | -0.036; 0.003 | 0.102 |
|  | Area Deprivation Index | 0.004 | 0.000 | 0.003; 0.005 | <0.001* |
|  | Sex | -0.057 | 0.013 | -0.081; -0.032 | <0.001* |
|  | CBCL- Exter | 0.005 | 0.001 | 0.003; 0.007 | <0.001* |
| Disorganized speech  - Distressing items score | Birth status | 0.008 | 0.034 | -0.059; 0.075 | 0.814 |
|  | Age | -0.006 | 0.006 | -0.019; 0.007 | 0.370 |
|  | Area Deprivation Index | 0.001 | 0.000 | 0.000; 0.001 | <0.001* |
|  | Sex | -0.023 | 0.008 | -0.039; -0.007 | 0.005* |
|  | CBCL- Exter | 0.003 | 0.001 | 0.002; 0.004 | <0.001* |
| Disorganized speech  - Distress severity score | Birth status | 0.024 | 0.091 | -0.154; 0.202 | 0.791 |
|  | Age | -0.013 | 0.017 | -0.046; 0.021 | 0.463 |
|  | Area Deprivation Index | 0.003 | 0.001 | 0.002; 0.004 | <0.001* |
|  | Sex | -0.044 | 0.022 | -0.087; -0.002 | 0.041* |
|  | CBCL- Exter | 0.010 | 0.002 | 0.007; 0.014 | <0.001* |

CBCL Child Behaviour Checklist; CI Confidence Interval; SE Standard Error. Birth status was coded as 0 = full-term birth and 1 = very preterm birth. Positive coefficients for Birth status therefore indicate higher symptom scores in the very preterm group relative to the full-term group

### **eResults 13**. Adjusted regression models of psychotic-like experiences in childhood in the ABCD cohort, additionally adjusted for Cognitive Ability and CBCL – Total problems score

| Outcome | Predictor | ß | SE | 95% CI | *p* value |
| --- | --- | --- | --- | --- | --- |
| Total PLEs  - Total score | Birth status | -0.252 | 0.297 | -0.833; 0.330 | 0.396 |
|  | Age | -0.271 | 0.054 | -0.377; -0.165 | <0.001* |
|  | Area Deprivation Index | 0.021 | 0.002 | 0.017; 0.025 | <0.001* |
|  | Sex | -0.249 | 0.069 | -0.384; -0.114 | <0.001* |
|  | CBCL- Total | -0.060 | 0.012 | -0.084; -0.037 | <0.001* |
|  | Cognitive Ability | 0.023 | 0.002 | 0.019; 0.027 | <0.001* |
| Total PLEs  - Distressing items score | Birth status | 0.056 | 0.191 | -0.319; 0.430 | 0.771 |
|  | Age | -0.206 | 0.035 | -0.275; -0.137 | <0.001* |
|  | Area Deprivation Index | 0.011 | 0.001 | 0.008; 0.014 | <0.001* |
|  | Sex | -0.042 | 0.045 | -0.129; 0.046 | 0.349 |
|  | CBCL- Total | -0.044 | 0.008 | -0.059; -0.029 | <0.001* |
|  | Cognitive Ability | 0.015 | 0.001 | 0.012; 0.017 | <0.001* |
| Total PLEs  - Distress severity score | Birth status | 0.259 | 0.619 | -0.953; 1.472 | 0.675 |
|  | Age | -0.638 | 0.115 | -0.863; -0.414 | <0.001* |
|  | Area Deprivation Index | 0.038 | 0.005 | 0.029; 0.048 | <0.001* |
|  | Sex | 0.062 | 0.145 | -0.223; 0.346 | 0.671 |
|  | CBCL- Total | -0.163 | 0.025 | -0.213; -0.114 | <0.001* |
|  | Cognitive Ability | 0.048 | 0.004 | 0.040; 0.056 | <0.001* |
| Unusual thought content  - Total score | Birth status | -0.154 | 0.152 | -0.452; 0.143 | 0.309 |
|  | Age | -0.124 | 0.028 | -0.179; -0.069 | <0.001* |
|  | Area Deprivation Index | 0.009 | 0.001 | 0.007; 0.012 | <0.001* |
|  | Sex | -0.031 | 0.036 | -0.101; 0.039 | 0.381 |
|  | CBCL- Total | -0.025 | 0.006 | -0.037; -0.012 | <0.001* |
|  | Cognitive Ability | 0.012 | 0.001 | 0.010; 0.014 | <0.001* |
| Unusual thought content  - Distressing items score | Birth status | 0.027 | 0.100 | -0.169; 0.223 | 0.786 |
|  | Age | -0.097 | 0.019 | -0.133; -0.060 | <0.001* |
|  | Area Deprivation Index | 0.005 | 0.001 | 0.003; 0.006 | <0.001* |
|  | Sex | 0.043 | 0.024 | -0.003; 0.090 | 0.065 |
|  | CBCL- Total | -0.019 | 0.004 | -0.027; -0.011 | <0.001* |
|  | Cognitive Ability | 0.007 | 0.001 | 0.006; 0.008 | <0.001* |
| Unusual thought content  - Distress severity score | Birth status | 0.080 | 0.336 | -0.578; 0.739 | 0.811 |
|  | Age | -0.323 | 0.062 | -0.445; -0.201 | <0.001* |
|  | Area Deprivation Index | 0.018 | 0.003 | 0.013; 0.023 | <0.001* |
|  | Sex | 0.237 | 0.079 | 0.082; 0.391 | 0.003* |
|  | CBCL- Total | -0.075 | 0.014 | -0.101; -0.048 | <0.001* |
|  | Cognitive Ability | 0.024 | 0.002 | 0.020; 0.029 | <0.001* |
| Perceptual abnormalities  - Total score | Birth status | -0.085 | 0.138 | -0.355; 0.185 | 0.536 |
|  | Age | -0.129 | 0.025 | -0.178; -0.079 | <0.001* |
|  | Area Deprivation Index | 0.008 | 0.001 | 0.006; 0.010 | <0.001* |
|  | Sex | -0.162 | 0.032 | -0.225; -0.099 | <0.001* |
|  | CBCL- Total | -0.022 | 0.006 | -0.033; -0.012 | <0.001* |
|  | Cognitive Ability | 0.010 | 0.001 | 0.008; 0.012 | <0.001* |
| Perceptual abnormalities  - Distressing items score | Birth status | 0.019 | 0.093 | -0.163; 0.201 | 0.841 |
|  | Age | -0.103 | 0.017 | -0.136; -0.069 | <0.001* |
|  | Area Deprivation Index | 0.005 | 0.001 | 0.004; 0.007 | <0.001* |
|  | Sex | -0.061 | 0.022 | -0.104; -0.019 | 0.005* |
|  | CBCL- Total | -0.020 | 0.004 | -0.027; -0.013 | <0.001* |
|  | Cognitive Ability | 0.006 | 0.001 | 0.005; 0.008 | <0.001* |
| Perceptual abnormalities  - Distress severity score | Birth status | 0.160 | 0.288 | -0.405; 0.725 | 0.579 |
|  | Age | -0.299 | 0.054 | -0.405; -0.194 | <0.001* |
|  | Area Deprivation Index | 0.018 | 0.002 | 0.014; 0.022 | <0.001* |
|  | Sex | -0.130 | 0.068 | -0.264; 0.003 | 0.055 |
|  | CBCL- Total | -0.072 | 0.012 | -0.095; -0.049 | <0.001* |
|  | Cognitive Ability | 0.020 | 0.002 | 0.016; 0.024 | <0.001* |
| Disorganized speech  - Total score | Birth status | -0.012 | 0.054 | -0.118; 0.093 | 0.819 |
|  | Age | -0.019 | 0.010 | -0.039; 0.001 | 0.063 |
|  | Area Deprivation Index | 0.003 | 0.000 | 0.003; 0.004 | <0.001* |
|  | Sex | -0.053 | 0.013 | -0.078; -0.028 | <0.001* |
|  | CBCL- Total | -0.014 | 0.002 | -0.018; -0.010 | <0.001* |
|  | Cognitive Ability | 0.002 | 0.000 | 0.001; 0.002 | <0.001* |
| Disorganized speech  - Distressing items score | Birth status | 0.008 | 0.034 | -0.059; 0.076 | 0.806 |
|  | Age | -0.006 | 0.007 | -0.019; 0.007 | 0.355 |
|  | Area Deprivation Index | 0.001 | 0.000 | 0.000; 0.001 | 0.008* |
|  | Sex | -0.022 | 0.008 | -0.038; -0.006 | 0.008* |
|  | CBCL- Total | -0.006 | 0.001 | -0.008; -0.003 | <0.001* |
|  | Cognitive Ability | 0.001 | 0.000 | 0.001; 0.002 | <0.001* |
| Disorganized speech  - Distress severity score | Birth status | 0.019 | 0.091 | -0.160; 0.198 | 0.836 |
|  | Age | -0.013 | 0.017 | -0.047; 0.021 | 0.467 |
|  | Area Deprivation Index | 0.002 | 0.001 | 0.001; 0.004 | 0.001* |
|  | Sex | -0.041 | 0.022 | -0.083; 0.002 | 0.063 |
|  | CBCL- Total | -0.018 | 0.004 | -0.025; -0.010 | <0.001* |
|  | Cognitive Ability | 0.004 | 0.001 | 0.003; 0.005 | <0.001* |

CBCL Child Behaviour Checklist; CI Confidence Interval; SE Standard Error. Birth status was coded as 0 = full-term birth and 1 = very preterm birth. Positive coefficients for Birth status therefore indicate higher symptom scores in the very preterm group relative to the full-term group

### **eResults 14**. Adjusted regression models of psychotic-like experiences in childhood in the ABCD cohort, additionally adjusted for Cognitive Ability and CBCL – Internalizing problems score

| Outcome | Predictor | ß | SE | 95% CI | *p* value |
| --- | --- | --- | --- | --- | --- |
| Total PLEs  - Total score | Birth status | -0.298 | 0.298 | -0.882; 0.286 | 0.317 |
|  | Age | -0.282 | 0.054 | -0.388; -0.175 | <0.001* |
|  | Area Deprivation Index | 0.022 | 0.002 | 0.018; 0.027 | <0.001* |
|  | Sex | -0.330 | 0.069 | -0.465; -0.195 | <0.001* |
|  | CBCL- Intern | -0.068 | 0.012 | -0.092; -0.045 | <0.001* |
|  | Cognitive Ability | 0.045 | 0.006 | 0.033; 0.058 | <0.001* |
| Total PLEs  - Distressing items score | Birth status | 0.026 | 0.192 | -0.350; 0.402 | 0.894 |
|  | Age | -0.214 | 0.035 | -0.283; -0.144 | <0.001* |
|  | Area Deprivation Index | 0.012 | 0.001 | 0.009; 0.015 | <0.001* |
|  | Sex | -0.093 | 0.045 | -0.180; -0.005 | 0.038* |
|  | CBCL- Intern | -0.049 | 0.008 | -0.064; -0.034 | <0.001* |
|  | Cognitive Ability | 0.030 | 0.004 | 0.022; 0.038 | <0.001* |
| Total PLEs  - Distress severity score | Birth status | 0.162 | 0.621 | -1.056; 1.380 | 0.794 |
|  | Age | -0.661 | 0.115 | -0.887; -0.435 | <0.001* |
|  | Area Deprivation Index | 0.041 | 0.005 | 0.031; 0.050 | <0.001* |
|  | Sex | -0.105 | 0.145 | -0.389; 0.179 | 0.469 |
|  | CBCL- Intern | -0.180 | 0.025 | -0.230; -0.131 | <0.001* |
|  | Cognitive Ability | 0.094 | 0.013 | 0.068; 0.121 | <0.001* |
| Unusual thought content  - Total score | Birth status | -0.178 | 0.153 | -0.477; 0.121 | 0.243 |
|  | Age | -0.130 | 0.028 | -0.185; -0.074 | <0.001* |
|  | Area Deprivation Index | 0.010 | 0.001 | 0.008; 0.012 | <0.001* |
|  | Sex | -0.072 | 0.035 | -0.142; -0.003 | 0.041* |
|  | CBCL- Intern | -0.029 | 0.006 | -0.041; -0.017 | <0.001* |
|  | Cognitive Ability | 0.023 | 0.003 | 0.016; 0.029 | <0.001* |
| Unusual thought content  - Distressing items score | Birth status | 0.013 | 0.100 | -0.184; 0.210 | 0.900 |
|  | Age | -0.100 | 0.019 | -0.137; -0.064 | <0.001* |
|  | Area Deprivation Index | 0.005 | 0.001 | 0.004; 0.007 | <0.001* |
|  | Sex | 0.019 | 0.023 | -0.027; 0.065 | 0.426 |
|  | CBCL- Intern | -0.021 | 0.004 | -0.029; -0.013 | <0.001* |
|  | Cognitive Ability | 0.014 | 0.002 | 0.010; 0.018 | <0.001* |
| Unusual thought content  - Distress severity score | Birth status | 0.031 | 0.337 | -0.630; 0.692 | 0.926 |
|  | Age | -0.335 | 0.062 | -0.457; -0.212 | <0.001* |
|  | Area Deprivation Index | 0.019 | 0.003 | 0.014; 0.024 | <0.001* |
|  | Sex | 0.152 | 0.079 | -0.002; 0.307 | 0.053 |
|  | CBCL- Intern | -0.083 | 0.014 | -0.110; -0.056 | <0.001* |
|  | Cognitive Ability | 0.048 | 0.007 | 0.034; 0.063 | <0.001* |
| Perceptual abnormalities  - Total score | Birth status | -0.105 | 0.138 | -0.376; 0.166 | 0.447 |
|  | Age | -0.134 | 0.025 | -0.183; -0.084 | <0.001* |
|  | Area Deprivation Index | 0.008 | 0.001 | 0.006; 0.010 | <0.001* |
|  | Sex | -0.196 | 0.032 | -0.259; -0.133 | <0.001* |
|  | CBCL- Intern | -0.026 | 0.006 | -0.037; -0.015 | <0.001* |
|  | Cognitive Ability | 0.021 | 0.003 | 0.015; 0.027 | <0.001* |
| Perceptual abnormalities  - Distressing items score | Birth status | 0.006 | 0.093 | -0.177; 0.188 | 0.952 |
|  | Age | -0.106 | 0.017 | -0.140; -0.072 | <0.001* |
|  | Area Deprivation Index | 0.006 | 0.001 | 0.004; 0.007 | <0.001* |
|  | Sex | -0.083 | 0.022 | -0.126; -0.041 | <0.001* |
|  | CBCL- Intern | -0.022 | 0.004 | -0.030; -0.015 | <0.001* |
|  | Cognitive Ability | 0.014 | 0.002 | 0.010; 0.018 | <0.001* |
| Perceptual abnormalities  - Distress severity score | Birth status | 0.119 | 0.289 | -0.447; 0.686 | 0.680 |
|  | Age | -0.309 | 0.054 | -0.415; -0.203 | <0.001* |
|  | Area Deprivation Index | 0.019 | 0.002 | 0.015; 0.023 | <0.001* |
|  | Sex | -0.200 | 0.068 | -0.333; -0.066 | 0.003* |
|  | CBCL- Intern | -0.079 | 0.012 | -0.102; -0.056 | <0.001* |
|  | Cognitive Ability | 0.039 | 0.006 | 0.027; 0.052 | <0.001* |
| Disorganized speech  - Total score | Birth status | -0.015 | 0.054 | -0.121; 0.090 | 0.776 |
|  | Age | -0.019 | 0.010 | -0.039; 0.000 | 0.056 |
|  | Area Deprivation Index | 0.004 | 0.000 | 0.003; 0.004 | <0.001* |
|  | Sex | -0.059 | 0.013 | -0.084; -0.034 | <0.001* |
|  | CBCL- Intern | -0.015 | 0.002 | -0.019; -0.010 | <0.001* |
|  | Cognitive Ability | 0.002 | 0.001 | -0.000; 0.004 | 0.080 |
| Disorganized speech  - Distressing items score | Birth status | 0.006 | 0.034 | -0.061; 0.073 | 0.862 |
|  | Age | -0.007 | 0.007 | -0.019; 0.006 | 0.314 |
|  | Area Deprivation Index | 0.001 | 0.000 | 0.000; 0.001 | 0.004* |
|  | Sex | -0.026 | 0.008 | -0.042; -0.010 | 0.002* |
|  | CBCL- Intern | -0.006 | 0.001 | -0.009; -0.003 | <0.001* |
|  | Cognitive Ability | 0.002 | 0.001 | 0.001; 0.004 | 0.003* |
| Disorganized speech  - Distress severity score | Birth status | 0.011 | 0.091 | -0.168; 0.190 | 0.902 |
|  | Age | -0.014 | 0.017 | -0.048; 0.020 | 0.409 |
|  | Area Deprivation Index | 0.002 | 0.001 | 0.001; 0.004 | <0.001* |
|  | Sex | -0.054 | 0.022 | -0.096; -0.011 | 0.014* |
|  | CBCL- Intern | -0.019 | 0.004 | -0.027; -0.012 | <0.001* |
|  | Cognitive Ability | 0.007 | 0.002 | 0.003; 0.011 | <0.001* |

CBCL Child Behaviour Checklist; CI Confidence Interval; SE Standard Error. Birth status was coded as 0 = full-term birth and 1 = very preterm birth. Positive coefficients for Birth status therefore indicate higher symptom scores in the very preterm group relative to the full-term group

### **eResults 15**. Adjusted regression models of psychotic-like experiences in childhood in the ABCD cohort, additionally adjusted for Cognitive Ability and CBCL – Externalizing problems score

| Outcome | Predictor | ß | SE | 95% CI | *p* value |
| --- | --- | --- | --- | --- | --- |
| Total PLEs  - Total score | Birth status | -0.239 | 0.297 | -0.822; 0.344 | 0.421 |
|  | Age | -0.267 | 0.054 | -0.373; -0.160 | <0.001* |
|  | Area Deprivation Index | 0.021 | 0.002 | 0.017; 0.026 | <0.001* |
|  | Sex | -0.257 | 0.069 | -0.393; -0.122 | <0.001* |
|  | CBCL- Extern | -0.060 | 0.012 | -0.084; -0.037 | <0.001* |
|  | Cognitive Ability | 0.057 | 0.006 | 0.045; 0.069 | <0.001* |
| Total PLEs  - Distressing items score | Birth status | 0.063 | 0.192 | -0.312; 0.439 | 0.741 |
|  | Age | -0.204 | 0.035 | -0.273; -0.135 | <0.001* |
|  | Area Deprivation Index | 0.011 | 0.001 | 0.009; 0.014 | <0.001* |
|  | Sex | -0.047 | 0.045 | -0.135; 0.041 | 0.296 |
|  | CBCL- Extern | -0.044 | 0.008 | -0.059; -0.029 | <0.001* |
|  | Cognitive Ability | 0.036 | 0.004 | 0.028; 0.044 | <0.001* |
| Total PLEs  - Distress severity score | Birth status | 0.287 | 0.621 | -0.929; 1.504 | 0.643 |
|  | Age | -0.630 | 0.115 | -0.855; -0.405 | <0.001* |
|  | Area Deprivation Index | 0.039 | 0.005 | 0.030; 0.048 | <0.001* |
|  | Sex | 0.050 | 0.146 | -0.235; 0.336 | 0.729 |
|  | CBCL- Extern | -0.163 | 0.025 | -0.213; -0.114 | <0.001* |
|  | Cognitive Ability | 0.121 | 0.013 | 0.096; 0.147 | <0.001* |
| Unusual thought content  - Total score | Birth status | -0.148 | 0.152 | -0.446; 0.151 | 0.332 |
|  | Age | -0.122 | 0.028 | -0.177; -0.067 | <0.001* |
|  | Area Deprivation Index | 0.010 | 0.001 | 0.007; 0.012 | <0.001* |
|  | Sex | -0.035 | 0.036 | -0.104; 0.035 | 0.332 |
|  | CBCL- Extern | -0.025 | 0.006 | -0.037; -0.013 | <0.001* |
|  | Cognitive Ability | 0.029 | 0.003 | 0.023; 0.036 | <0.001* |
| Unusual thought content  - Distressing items score | Birth status | 0.031 | 0.100 | -0.166; 0.228 | 0.757 |
|  | Age | -0.096 | 0.019 | -0.132; -0.059 | <0.001* |
|  | Area Deprivation Index | 0.005 | 0.001 | 0.003; 0.006 | <0.001* |
|  | Sex | 0.042 | 0.024 | -0.005; 0.088 | 0.079 |
|  | CBCL- Extern | -0.019 | 0.004 | -0.027; -0.011 | <0.001* |
|  | Cognitive Ability | 0.018 | 0.002 | 0.014; 0.022 | <0.001* |
| Unusual thought content  - Distress severity score | Birth status | 0.095 | 0.337 | -0.566; 0.755 | 0.779 |
|  | Age | -0.319 | 0.062 | -0.442; -0.197 | <0.001* |
|  | Area Deprivation Index | 0.018 | 0.003 | 0.013; 0.023 | <0.001* |
|  | Sex | 0.231 | 0.079 | 0.076; 0.386 | 0.003* |
|  | CBCL- Extern | -0.074 | 0.014 | -0.101; -0.048 | <0.001* |
|  | Cognitive Ability | 0.062 | 0.007 | 0.048; 0.075 | <0.001* |
| Perceptual abnormalities  - Total score | Birth status | -0.081 | 0.138 | -0.351; 0.190 | 0.559 |
|  | Age | -0.127 | 0.025 | -0.177; -0.077 | <0.001* |
|  | Area Deprivation Index | 0.008 | 0.001 | 0.006; 0.010 | <0.001* |
|  | Sex | -0.166 | 0.032 | -0.230; -0.103 | <0.001* |
|  | CBCL- Extern | -0.023 | 0.006 | -0.034; -0.012 | <0.001* |
|  | Cognitive Ability | 0.023 | 0.003 | 0.017; 0.029 | <0.001* |
| Perceptual abnormalities  - Distressing items score | Birth status | 0.022 | 0.093 | -0.161; 0.204 | 0.817 |
|  | Age | -0.102 | 0.017 | -0.135; -0.068 | <0.001* |
|  | Area Deprivation Index | 0.005 | 0.001 | 0.004; 0.007 | <0.001* |
|  | Sex | -0.064 | 0.022 | -0.107; -0.021 | 0.003* |
|  | CBCL- Extern | -0.020 | 0.004 | -0.028; -0.013 | <0.001* |
|  | Cognitive Ability | 0.015 | 0.002 | 0.011; 0.019 | <0.001* |
| Perceptual abnormalities  - Distress severity score | Birth status | 0.171 | 0.289 | -0.395; 0.737 | 0.554 |
|  | Age | -0.296 | 0.054 | -0.402; -0.191 | <0.001* |
|  | Area Deprivation Index | 0.018 | 0.002 | 0.014; 0.022 | <0.001* |
|  | Sex | -0.135 | 0.068 | -0.269; -0.002 | 0.047* |
|  | CBCL- Extern | -0.072 | 0.012 | -0.095; -0.049 | <0.001* |
|  | Cognitive Ability | 0.050 | 0.006 | 0.038; 0.062 | <0.001* |
| Disorganized speech  - Total score | Birth status | -0.011 | 0.054 | -0.116; 0.094 | 0.835 |
|  | Age | -0.019 | 0.010 | -0.038; 0.001 | 0.067 |
|  | Area Deprivation Index | 0.003 | 0.000 | 0.003; 0.004 | <0.001* |
|  | Sex | -0.054 | 0.013 | -0.079; -0.029 | <0.001* |
|  | CBCL- Extern | -0.014 | 0.002 | -0.018; -0.010 | <0.001* |
|  | Cognitive Ability | 0.004 | 0.001 | 0.002; 0.007 | <0.001* |
| Disorganized speech  - Distressing items score | Birth status | 0.009 | 0.034 | -0.058; 0.076 | 0.795 |
|  | Age | -0.006 | 0.007 | -0.019; 0.007 | 0.366 |
|  | Area Deprivation Index | 0.001 | 0.000 | 0.000; 0.001 | 0.006* |
|  | Sex | -0.022 | 0.008 | -0.038; -0.006 | 0.007* |
|  | CBCL- Extern | -0.006 | 0.001 | -0.008; -0.003 | <0.001* |
|  | Cognitive Ability | 0.003 | 0.001 | 0.001; 0.004 | <0.001* |
| Disorganized speech  - Distress severity score | Birth status | 0.021 | 0.091 | -0.158; 0.200 | 0.817 |
|  | Age | -0.012 | 0.017 | -0.046; 0.022 | 0.485 |
|  | Area Deprivation Index | 0.002 | 0.001 | 0.001; 0.004 | <0.001* |
|  | Sex | -0.041 | 0.022 | -0.084; 0.002 | 0.059 |
|  | CBCL- Extern | -0.018 | 0.004 | -0.025; -0.010 | <0.001* |
|  | Cognitive Ability | 0.010 | 0.002 | 0.006; 0.013 | <0.001* |

CBCL Child Behaviour Checklist; CI Confidence Interval; SE Standard Error. Birth status was coded as 0 = full-term birth and 1 = very preterm birth. Positive coefficients for Birth status therefore indicate higher symptom scores in the very preterm group relative to the full-term group

### **eResults 16.** Pooled analyses across the BIPP and ABCD cohorts. Model adjusted for cohort, age, and sex

| Outcome | Predictor | ß | Robust SE (HC3) | 95% CI | *p* value |
| --- | --- | --- | --- | --- | --- |
| Total PLEs  - Total score | Cohort (ABCD vs BIPP) | -1.253 | 0.242 | -1.727; -0.778 | <0.001* |
|  | Age | -0.286 | 0.050 | -0.384: -0.187 | <0.001* |
|  | Sex | -0.289 | 0.068 | -0.422; -0.156 | <0.001* |
| Total PLEs  - Distressing items score | Cohort (ABCD vs BIPP) | -0.721 | 0.169 | -1.053; -0.389 | <0.001* |
|  | Age | -0.186 | 0.033 | -0.252; -0.121 | <0.001* |
|  | Sex | -0.066 | 0.043 | -0.150; 0.019 | 0.130 |
| Total PLEs  - Distress severity score | Cohort (ABCD vs BIPP) | -2.315 | 0.559 | -3.411; -1.218 | <0.001* |
|  | Age | -0.565 | 0.113 | -0.786; -0.344 | <0.001* |
|  | Sex | -0.043 | 0.141 | -0.319; 0.233 | 0.760 |
| Unusual thought content  - Total score | Cohort (ABCD vs BIPP) | -0.502 | 0.119 | -0.736; -0.269 | <0.001* |
|  | Age | -0.133 | 0.025 | -0.183; -0.084 | <0.001* |
|  | Sex | -0.043 | 0.035 | -0.111; 0.025 | 0.213 |
| Unusual thought content  - Distressing items score | Cohort (ABCD vs BIPP) | -0.229 | 0.082 | -0.390; -0.068 | 0.005* |
|  | Age | -0.086 | 0.017 | -0.119; -0.053 | <0.001* |
|  | Sex | 0.032 | 0.023 | -0.013; 0.076 | 0.162 |
| Unusual thought content  - Distress severity score | Cohort (ABCD vs BIPP) | -0.800 | 0.281 | -1.351; -0.250 | 0.004* |
|  | Age | -0.283 | 0.057 | -0.394; -0.172 | <0.001* |
|  | Sex | 0.176 | 0.076 | 0.027; 0.324 | 0.021* |
| Perceptual abnormalities  - Total score | Cohort (ABCD vs BIPP) | -0.417 | 0.106 | -0.624; -0.209 | <0.001* |
|  | Age | -0.127 | 0.023 | -0.172; -0.083 | <0.001* |
|  | Sex | -0.194 | 0.031 | -0.255; -0.133 | <0.001* |
| Perceptual abnormalities  - Distressing items score | Cohort (ABCD vs BIPP) | -0.266 | 0.075 | -0.414; -0.119 | <0.001* |
|  | Age | -0.089 | 0.016 | -0.121; -0.058 | <0.001* |
|  | Sex | -0.080 | 0.021 | -0.121; -0.039 | <0.001* |
| Perceptual abnormalities  - Distress severity score | Cohort (ABCD vs BIPP) | -0.785 | 0.227 | -1.230; -0.341 | <0.001* |
|  | Age | -0.258 | 0.051 | -0.357; -0.158 | <0.001* |
|  | Sex | -0.189 | 0.065 | -0.317; -0.061 | 0.004* |
| Disorganized speech  - Total score | Cohort (ABCD vs BIPP) | -0.171 | 0.043 | -0.255; -0.088 | <0.001* |
|  | Age | -0.020 | 0.009 | -0.038; -0.002 | 0.029* |
|  | Sex | -0.052 | 0.012 | -0.076; -0.027 | <0.001* |
| Disorganized speech  - Distressing items score | Cohort (ABCD vs BIPP) | -0.108 | 0.032 | -0.171; -0.046 | <0.001* |
|  | Age | -0.007 | 0.006 | -0.019; 0.005 | 0.243 |
|  | Sex | -0.018 | 0.008 | -0.033; -0.002 | 0.024* |
| Disorganized speech  - Distress severity score | Cohort (ABCD vs BIPP) | -0.354 | 0.099 | -0.548; -0.159 | <0.001* |
|  | Age | -0.017 | 0.018 | -0.052; 0.017 | 0.329 |
|  | Sex | -0.039 | 0.021 | -0.075; 0.008 | 0.111 |

CI Confidence Interval; HC3 Heteroscedasticity-consistent covariance matrix estimator; SE Standard Error. Birth status was coded as 0 = full-term birth and 1 = very preterm birth. Positive coefficients for Birth status therefore indicate higher symptom scores in the very preterm group relative to the full-term group.

### **eResults 17.** Pooled analyses across the BIPP and ABCD cohorts. Model adjusted for cohort, age, sex, and birth status

| Outcome | Predictor | ß | Robust SE (HC3) | 95% CI | *p* value |
| --- | --- | --- | --- | --- | --- |
| Total PLEs  - Total score | Cohort (ABCD vs BIPP) | -0.869 | 0.277 | -1.412; -0.327 | 0.002* |
|  | Age | -0.298 | 0.051 | -0.397; -0.199 | <0.001* |
|  | Sex | -0.289 | 0.068 | -0.422; -0.156 | <0.001* |
|  | Birth status | 0.545 | 0.229 | 0.095; 0.994 | 0.018* |
| Total PLEs  - Distressing items score | Cohort (ABCD vs BIPP) | -0.419 | 0.194 | -0.800; -0.038 | 0.031* |
|  | Age | -0.196 | 0.033 | -0.262; -0.131 | <0.001* |
|  | Sex | -0.065 | 0.043 | -0.150; 0.019 | 0.131 |
|  | Birth status | 0.429 | 0.155 | 0.125; 0.733 | 0.006* |
| Total PLEs  - Distress severity score | Cohort (ABCD vs BIPP) | -1.331 | 0.613 | -2.532; -0.130 | 0.030* |
|  | Age | -0.598 | 0.113 | -0.819; -0.376 | <0.001* |
|  | Sex | -0.043 | 0.141 | -0.319; 0.233 | 0.761 |
|  | Birth status | 1.398 | 0.504 | 0.409; 2.386 | 0.006* |
| Unusual thought content  - Total score | Cohort (ABCD vs BIPP) | -0.363 | 0.143 | -0.644; -0.083 | 0.011* |
|  | Age | -0.138 | 0.026 | -0.188; -0.088 | <0.001* |
|  | Sex | -0.043 | 0.035 | -0.111; 0.025 | 0.214 |
|  | Birth status | 0.197 | 0.120 | -0.038; 0.433 | 0.101 |
| Unusual thought content  - Distressing items score | Cohort (ABCD vs BIPP) | -0.090 | 0.097 | -0.281; 0.100 | 0.353 |
|  | Age | -0.091 | 0.017 | -0.124; -0.058 | <0.001* |
|  | Sex | 0.032 | 0.023 | -0.013; 0.076 | 0.162 |
|  | Birth status | 0.197 | 0.083 | 0.034; 0.359 | 0.018* |
| Unusual thought content  - Distress severity score | Cohort (ABCD vs BIPP) | -0.374 | 0.326 | -1.013; 0.264 | 0.250 |
|  | Age | -0.297 | 0.057 | -0.409; -0.186 | <0.001* |
|  | Sex | 0.176 | 0.076 | 0.027; 0.325 | 0.021* |
|  | Birth status | 0.605 | 0.284 | 0.049; 1.161 | 0.033* |
| Perceptual abnormalities  - Total score | Cohort (ABCD vs BIPP) | -0.210 | 0.119 | -0.443; 0.023 | 0.077 |
|  | Age | -0.134 | 0.023 | -0.179; -0.089 | <0.001* |
|  | Sex | -0.194 | 0.031 | -0.255; -0.133 | <0.001* |
|  | Birth status | 0.294 | 0.106 | 0.087; 0.500 | 0.005* |
| Perceptual abnormalities  - Distressing items score | Cohort (ABCD vs BIPP) | -0.110 | 0.085 | -0.278; 0.057 | 0.198 |
|  | Age | -0.095 | 0.016 | -0.126; -0.063 | <0.001* |
|  | Sex | -0.080 | 0.021 | -0.121; -0.039 | <0.001* |
|  | Birth status | 0.222 | 0.073 | 0.078; 0.366 | 0.002* |
| Perceptual abnormalities  - Distress severity score | Cohort (ABCD vs BIPP) | -0.275 | 0.244 | -0.754; 0.204 | 0.260 |
|  | Age | -0.275 | 0.051 | -0.375; -0.175 | <0.001* |
|  | Sex | -0.189 | 0.065 | -0.316; -0.051 | 0.004* |
|  | Birth status | 0.725 | 0.216 | 0.301; 1.149 | <0.001* |
| Disorganized speech  - Total score | Cohort (ABCD vs BIPP) | -0.137 | 0.051 | -0.238; -0.036 | 0.008* |
|  | Age | -0.021 | 0.009 | -0.039; -0.003 | 0.022* |
|  | Sex | -0.052 | 0.012 | -0.076; -0.027 | <0.001* |
|  | Birth status | 0.049 | 0.044 | -0.037; 0.134 | 0.267 |
| Disorganized speech  - Distressing items score | Cohort (ABCD vs BIPP) | -0.096 | 0.039 | -0.173; -0.020 | 0.014* |
|  | Age | -0.008 | 0.006 | -0.020; 0.004 | 0.218 |
|  | Sex | -0.018 | 0.008 | -0.033; -0.002 | 0.024* |
|  | Birth status | 0.017 | 0.031 | -0.043; 0.077 | 0.581 |
| Disorganized speech  - Distress severity score | Cohort (ABCD vs BIPP) | -0.275 | 0.112 | -0.494; -0.056 | 0.014* |
|  | Age | -0.020 | 0.018 | -0.054; 0.015 | 0.260 |
|  | Sex | -0.034 | 0.021 | -0.075; 0.008 | 0.111 |
|  | Birth status | 0.111 | 0.089 | -0.063; 0.285 | 0.212 |

CI Confidence Interval; HC3 Heteroscedasticity-consistent covariance matrix estimator; SE Standard Error. Birth status was coded as 0 = full-term birth and 1 = very preterm birth. Positive coefficients for Birth status therefore indicate higher symptom scores in the very preterm group relative to the full-term group.
